# Confidence Analysis of DEER Data and its Structural Interpretation with Ensemble-Biased Metadynamics

**DOI:** 10.1101/299941

**Authors:** Eric J. Hustedt, Fabrizio Marinelli, Richard A. Stein, José D. Faraldo-Gómez, Hassane S. Mchaourab

**Author notes:** EJH and FM contributed equally. Correspondence should be addressed to: JDFG, HSM.

## Abstract

Given its ability to measure multicomponent distance distributions between electron-spin probes, Double Electron-Electron Resonance spectroscopy (DEER) has become a leading technique to assess the structural dynamics of biomolecules. However, methodologies to evaluate the statistical error of these distributions are not standard, often hampering a rigorous interpretation of the experimental results. Distance distributions are often determined from the experimental DEER data through a mathematical method known as Tikhonov regularization, but this approach makes rigorous error estimates difficult. Here, we build upon an alternative model-based approach in which the distance probability distribution is represented as a sum of Gaussian components and use propagation of errors to calculate an associated confidence band. Our approach considers all sources of uncertainty, including the experimental noise, the uncertainty in the fitted background signal, and the limited time-span of the data collection. The resulting confidence band reveals the most and least reliable features of the probability distribution, thereby informing the structural interpretation of DEER experiments. To facilitate this interpretation, we also generalize the molecular-simulation method known as Ensemble-Biased Metadynamics. This method, originally designed to generate maximum-entropy structural ensembles consistent with one or more probability distributions, now also accounts for the uncertainty in those target distributions, exactly as dictated by their confidence bands. After careful benchmarks, we demonstrate the proposed techniques using DEER results from spin-labeled T4 lysozyme.

## INTRODUCTION

Double Electron-Electron Resonance (**DEER**) spectroscopy is a pulsed electron-spin resonance technique that is widely used to measure long-range distances between paramagnetic species, typically extrinsic probes introduced into biological macromolecules by some form of site-directed spin labeling (1-3). The main advantage of DEER lies in its ability to go beyond measuring the average distance between labels and resolve complex distance distributions that depend on both the rotameric states of the spin labels and also on differences in backbone structure of the protein or other biomolecule. It is this sensitivity to distinct backbone conformations that allows DEER experiments to give unique insights into the structure and functional dynamics of the protein under study (4).

The translation of an experimental time-domain DEER signal, *D*(*t*), into a distance distribution, *P*(*R*), is, however, an ill-posed mathematical problem in that small variations in *D*(*t*) can lead to large variations in the *P*(*R*) obtained. To address this issue, approaches to the analysis of DEER data impose some degree of smoothness on *P*(*R*), either by adding an adjustable smoothness factor to the fit criterion via Tikhonov regularization (**TR**) (5-8), or by assuming some smooth functional form, such as a sum of Gaussian components, to model *P*(*R*) (9-12).

The experimental DEER time-domain signal is the product of a factor arising from the dipolar interactions between the small number of spins (typically two) within a labeled molecule, *D*_O_(*t*), and a background signal, *D*_B_(*t*), arising from a large number of intermolecular dipolar interactions. Thus,

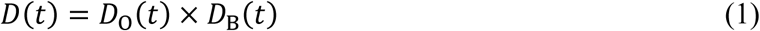

Properly accounting for the background signal is therefore necessary to determine the desired intramolecular distance distributions that are reflected in *D*_O_(*t*).

Application of the TR method requires that an estimate of the background factor be made *a priori* by fitting the latter portion of the time-domain signal *D*(*t*). This estimated background factor enables the determination of a background-corrected signal as an estimate of *D*_O_(*t*) that is then analyzed to give a distance distribution. Error estimations are typically made *a posteriori*, by assessing the effects of the background correction and of the experimental noise at an arbitrary statistical-significance level. More recently, a rigorous Bayesian approach has been developed within the TR framework for quantifying the uncertainty in *P*(*R*) due to the experimental noise and the uncertainty in the choice of the optimal regularization parameter (13). This Bayesian approach does not, as of yet, allow for an estimate in the uncertainty of *P*(*R*) due to the background correction.

An alternative model-based approach, in which *P*(*R*) is represented as a sum of smooth basis functions, e.g. Gaussian components, relies on the simultaneous determination of the best-fit parameters modeling both s*D*_B_(*t*) and *D*_O_(*t*) by a non-linear least-squares algorithm. The advantages of a model-based approach for the analysis of DEER data, as opposed to *a priori* background correction followed by TR, have been detailed previously (9, 10). One of these advantages is the ability to perform a rigorous error analysis on the various fit parameters including those used to define *P*(*R*). For multicomponent distance distributions, however, it can be difficult to appreciate how the parameter uncertainties affect the confidence in the resulting *P*(*R*).

In this work, a robust and computationally efficient algorithm is developed to quantify the uncertainty in *P*(*R*) in terms of a confidence band about the best-fit solution. This confidence band reflects the influence of both the noise in the measured data and the uncertainty in the estimate of the background correction, and can be calculated with no significant increase in computation time. The algorithm uses the method of propagation of errors, otherwise known as the delta method, to estimate the variance in a function, here *P*(*R*), of a set of random variables, here all the best-fit parameters for a given *D*(*t*) (14-16). We demonstrate the validity and robustness of the delta method as applied to the analysis of DEER data using different sets of simulated data. Then, we analyze experimental data from T4 lysozyme (**T4L**) using the new algorithm. Confidence bands obtained using the delta method quantify the reliability of each of the features of the distance distributions thus permitting an objective comparison of results from different experiments.

Once estimates of *P*(*R*) and its associated error have been obtained, the next step is to use these data for assessing the structural dynamics of a biomolecule. This is a non-trivial task since *P*(*R*) reflects both variations in the backbone structure and conformational flexibility of the spin labels. Even for a rigid protein, for example, different rotamers of MTSSL labels can result in ~10-Å wide distance distributions featuring multiple peaks (17-20). Molecular dynamics (MD) simulations are arguably the most rigorous approach to model this variability. Of particular value are advanced simulation approaches that implement a bias on the calculated trajectories to reproduce the experimentally determined *P*(*R*) while fulfilling the so-called maximum-entropy condition, i.e. when the bias applied is the minimum required (17, 18). To our knowledge, however, none of the biasing techniques of this kind considers explicitly the uncertainty of the target data, which effectively violates the minimum-information condition and can potentially lead to erroneous interpretations of the data. Here, we generalize one of these advanced simulation techniques, known as Ensemble-Biased Metadynamics (EBMetaD), so that it rigorously accounts for the uncertainty of the input data. Like the original EBMetaD, the method is based on an adaptive biasing algorithm that gradually develops a molecular ensemble consistent with the target distribution. However, the bias applied is the least required for the *P*(*R*) from the MD trajectories to be consistent with the confidence bands of the experimental results. The EBMetaD method can be applied to a probability distribution corresponding to any molecular variable, either obtained through any experimental method or postulated theoretically. This innovative simulation methodology is first benchmarked on a small-molecule system using a hypothetical distribution corresponding to a dihedral angle. Then, EBMetaD is applied to the abovementioned DEER data obtained for **T4L** to construct the corresponding structural ensembles in explicit water.

## METHODS

### Simulated and Experimental DEER signals

Simulated DEER data were generated using the program DEERsim version 2 running in Matlab R2017a (The Mathworks Inc., Natick MA) with artificial noise added in the form of normally-distributed random numbers with a given standard deviation. DEERsim is based on previously published algorithms for calculating DEER time-domain signals (9), and is freely available at https://lab.vanderbilt.edu/hustedt-lab/software. In some cases 10,000 replicate data sets were created from the same noiseless time trace and used to evaluate, via Monte Carlo simulations, the proposed methodologies for estimating the confidence in the fit parameters and in *P*(*R*). The experimental DEER signals for the 3 doubly-labeled mutants of T4 lysozyme (namely residues 62 and 109, 62 and 134 and 109 and 134) were taken from previously published work (21, 22).

### Analysis of DEER Data

The simulated and experimental DEER data were analyzed using the program DD version 6C running in Matlab R2017a as previously described (10) with modifications to allow for: 1) the calculation of Bayesian Information Criterion values; 2) the estimation of parameter uncertainties from the variance-covariance matrix; and 3) the calculation of a confidence band for the best-fit *P*(*R*) using the delta method. Details on each of these three new procedures are provided below. DD is freely available at https://lab.vanderbilt.edu/hustedt-lab/software.

Assuming an ideal three-dimensional solution, simulated DEER signals, *F*(*t*), are modeled according to

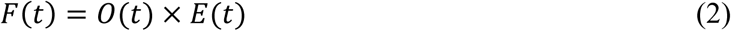

where *O*(*t*) is the calculated signal for the pair of spins within a molecule,

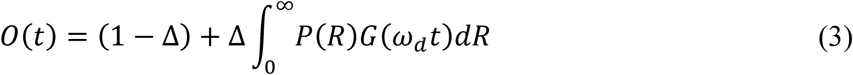

and *E*(*t*) is an exponential to account for the background intermolecular interactions.

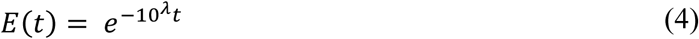

Here, Δ is the modulation-depth parameter, *λ* is a parameter governing the exponential background decay rate, *P*(*R*) is any probability distribution for the intramolecular inter-electron distance, and *G*(*ω_d_t*) is a kernel function defined previously (5, 13, 23),

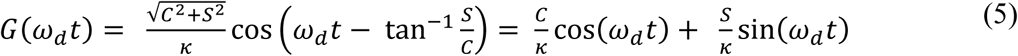

where *C* and *S* are the Fresnel cosine and sine integrals

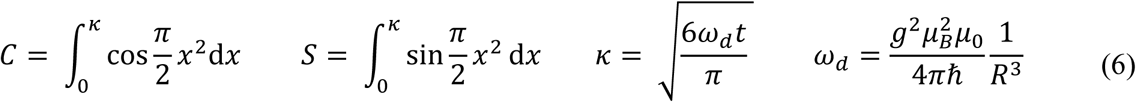

and the symbols in the equation for *ω_d_* represent the usual physical constants.

Our analysis of DEER data is based on the assumption that *P*(*R*) can be described by a sum of *n* Gaussian components:

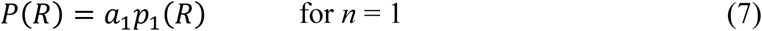

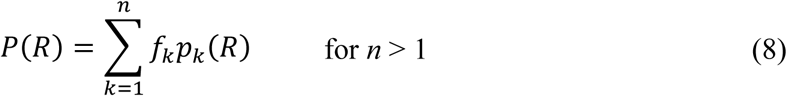

where *a*_1_ *≡* 1 and

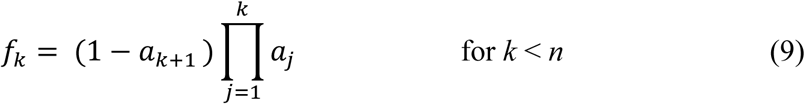

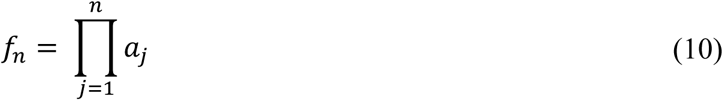

The Gaussian components are given by

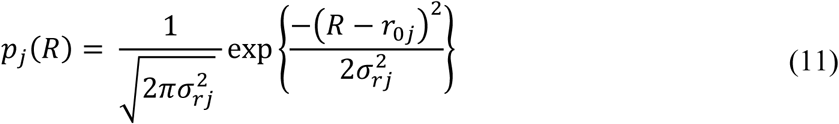

Alternative basis functions with non-Gaussian shapes may also be used (see Supplemental Methods). The use of the *a_j>_*_2_ as fit parameters guarantees that for any value of *n* and any set of 0≤ *a_j>_*_2_ ≤1 the resulting *P*(*R*) will be normalized. For a given number of components, *n*, the set of 3n-1 variables *r_0j_*, *σ_rj_*, and *a_j>_*_2_ define *P*(*R*). All of these variables together with Δ, *λ*, and a scale factor constitute the set of parameters, *β_l_*_=1,2,….*q*_ that need to be determined for a given experimental signal and a given model (i.e. a specific value of *n*). We define the best-fit values of these parameters as those that minimize the reduced chi-squared value:

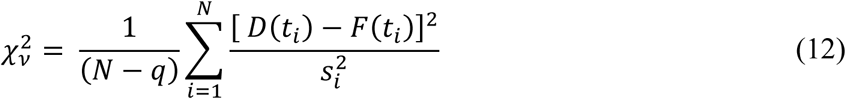

where *D*(*t*) is the experimental time-domain DEER data, *N* is the total number of data points, *q* is the total number of parameters considered in the fit, *s_i_* is the estimated noise level (standard deviation) of the *i*^th^ data point, and *t_i_* is the time value of the *i*^th^ data point. Here, the noise level is assumed to be uniform (*s_i_* = s) at a level estimated from the standard deviation of the imaginary component of the data.

For comparison, the simulated data were also analyzed using DeerAnalysis2016 (http://www.epr.ethz.ch/software/index) using an *a priori* background correction and TR (6). The zero time, phase correction, and initial start time for background fitting were determined automatically using the “!” button. When necessary, the regularization parameter was manually adjusted to match the corner of the *L* curve. The “Validation” tool was used to estimate the uncertainty in *P*(*R*) due to a range of starting times for background fitting with results pruned to eliminate those that increased the RMS deviation by more than 15% as recommended, although the statistical significance of this increase is unknown.

### Bayesian Information Criterion

Previously, the Akaike information criterion corrected for finite sample size (***AICc***) has been used to select the optimal model for a given experimental signal (10). Here, the closely related Bayesian Information Criterion (***BIC***) is used:

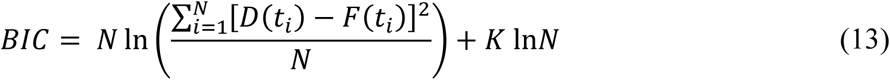

where *K* = *q* + 1. *BIC* can be used to select the optimum number of Gaussians describing *P*(*R*) that explain the data without overfitting (24). The optimal value of *n* is the one that results in the lowest *BIC*. *BIC* differs from *AICc* only in the second term of Equation 13. For typical values of *N* and *q* (*i.e. N* ≥ *85* and *q* ≥ 26), *BIC* will always increase faster than *AICc* with increasing q. Thus, *BIC* will favor the same model or a more parsimonious models, *i.e*. a lower value of n, and in our judgement is preferable. For a given model *j*, Δ*BICj* is given by

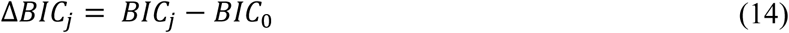

where *BIC*_0_ is the lowest *BIC* value obtained for a given data set. *AIC*c, *BIC*, and related criterion have also been recently evaluated by Edwards and Stoll as methods to determine the optimal regularization parameter for TR analysis of DEER data (25).

### Parameter Uncertainties

The methodology proposed herein aims to not only identify the best-fit values of the set of parameters *β_l=_*_1,2,….*q*_ but also their uncertainty. This uncertainty can be rigorously quantified for each parameter by calculating a series of 1-dimensional confidence intervals (26, 27) as described in detail elsewhere (10). Alternatively, under appropriate conditions the parameter uncertainties can be estimated from the standard errors determined from the covariance matrix *C* = *α*^−1^, where *α* is the curvature matrix whose elements are:

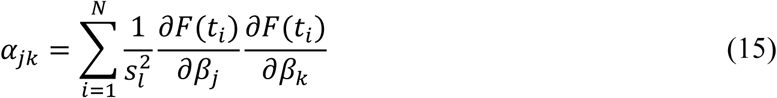

where the required partial derivatives are determined numerically via the forward difference method. The standard errors of each of the parameters *l*, *σ_l_*, are determined from the diagonal elements of *C*:

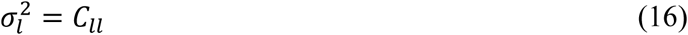

and the off-diagonal elements give the covariances between parameters. The fit parameters and their uncertainties are reported as:

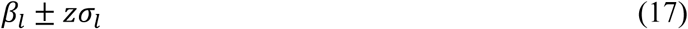

where *z* = 1, 2, or 3 depending on whether the confidence level desired is 1*σ* (68.3%), 2*σ* (95.4%), or 3*σ* (99.7%). In contrast to the calculation of confidence intervals, estimating the parameter uncertainties from the covariance matrix requires little additional computation. The validity of both approaches will be assessed below using a Monte Carlo approach.

### Confidence Bands

The confidence band for a *P*(*R*) is calculated from the full covariance matrix using the delta method (14-16). Given that the best-fit parameters are themselves random variables obtained from fitting a given data set, we have

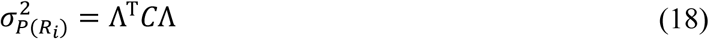

where Λ is a vector of the partial derivatives of *P*(*R*) at a particular distance *R_i_* with respect to all of the fit parameters *β_j_*_=1,2_*_,….q_*, i.e.

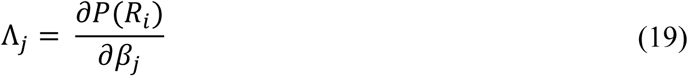

Here the partial derivatives of *P*(*R*) with respect to *a_j>_*_2_ are determined analytically, those with respect to *r_0j_* and *σ_rj_* are determined numerically, and those with respect to other parameters such as Δ, *λ*, and the scale factor are strictly zero. The confidence band for *P*(*R*) is then given by

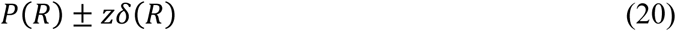

where

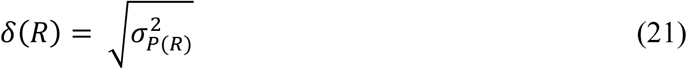

and *z* = 1, 2, or 3 depending on whether a band at the 1σ, 2σ, or 3σ confidence level is desired.

### Molecular Dynamics Simulations

All MD simulations were carried out with NAMD versions 2.9-2.12 (28). The force-field used was CHARMM27/CMAP (29, 30), augmented by a force field for the spin labels developed by Sezer et al. (31). The simulations were carried out at 298 K and 1 bar with a 2-fs time-step and periodic boundary conditions. Van der Waals and short-range electrostatic interactions were cut off at 12 Å; the PME method was used to calculate long-range electrostatic interactions. Two molecular systems are considered, butyramide in a cubic box with 1467 water molecules and T4L in a truncated-octahedral box with 12,013 water molecules and Cl^-^ counter ions to neutralize the total charge.

### Ensemble-Biased Metadynamics: Formalism

We introduce a generalization of the Ensemble-Biased Metadynamics (EBMetaD) technique (18) to take into account the confidence band on the distance distribution determined from experimental data. In MD simulations based on the EBMetaD method, a function of the atomic coordinates **X**, *ξ* = *ξ^f^*[**X**], is defined and a time-dependent biasing potential *V*(*ξ*,*t*) is added to the standard energy function to ensure that the ensemble of conformations explored during the simulation is consistent with a given target probability distribution, *ρ*(*ξ*). The biasing potential is gradually constructed as a sum of Gaussian functions of *ξ*, added at time intervals *τ* and centered on the instantaneous value of *ξ* (18):

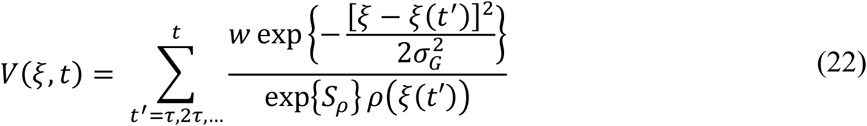

where *ξ*(*t*′) denotes the value of *ξ* at time *t*′, *σ_G_* is related to the resolution used to describe fluctuations of *ξ*, *w* is a scaling parameter of the Gaussians height and exp{*S_ρ_*} is the effective volume spanned by *ρ*(*ξ*) (i.e. *S_ρ_* = − ∫ *ρ*(*ξ*)ln[*ρ*(*ξ*)]*dξ* is the differential entropy of *ρ*(*ξ*). In the original application of the EBMetaD approach to DEER spectroscopy (18), the collective coordinate is the inter-label distance (ξ = *R*) and the target probability density is the DEER distance distribution (*ρ*(*ξ*)*=P*(*R*)), thereby assuming that the uncertainty on *P*(*R*) is negligible. Following the same notation, from here on we denote the experimental best-fit distribution as *ρ*(*ξ*), and its uncertainty is represented by *δ*(*ξ*).

To account for this uncertainty, the new EBMetaD approach targets not *ρ*(*ξ*), but *ρ*(*ξ*) + *δ*(*ξ*). More specifically, the desired simulated ensemble corresponds to a distribution that satisfies two requirements: (1) it is inside the experimental confidence band; and (2) it minimizes the amount of bias added to the standard energy function, i.e. it resembles the unbiased probability distribution of a conventional MD simulation as much as possible. In practice, this approach requires that the simulation be biased to sample an adaptive distribution denoted as *ρ*(*ξ*, *t*). This distribution varies as the simulation evolves, so as to ensure conditions (1) and (2) are ultimately fulfilled. An expression for *ρ*(*ξ*, *t*) can be derived either from an extended formulation of the maximum-entropy principle (32) that considers the experimental uncertainty (33, 34) or from a Bayesian approach (35):

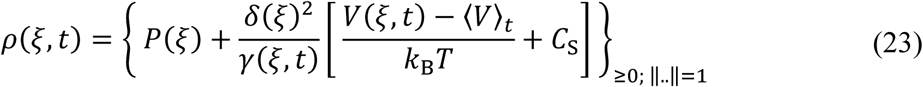

where *k*_B_ is the Boltzmann constant, *T* is the simulation temperature, and *C_S_* is a shift constant. Here, 〈*V*〉*_t_* denotes the average value of the biasing potential at time *t*, i.e. 〈*V*〉*_t_* = ∫ *ρ*(*ξ*, *t*) *Vρ*(*ξ*, *t*) *dξ*, which serves as an offset of the instantaneous biasing potential *V*(*ξ*, *t*). The term *γ*(*ξ*, *t*)is a scaling factor of *δ*(*ξ*), initially set to 1 and then updated during the simulation, as discussed below. The notation 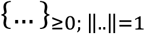 denotes a projection onto the probability simplex (36) and guarantees that *ρ*(*ξ*, *t*) is positive and normalized, i.e. ∫ *ρ*(*ξ*, *t*) *dξ* = 1. This normalization condition, in turn, sets the value of *C_S_*. That is,

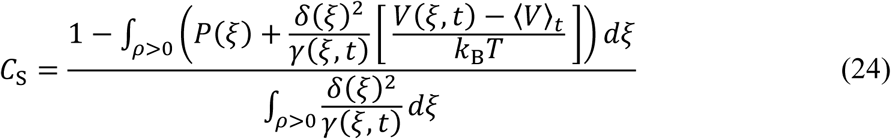

Note that in Eq. 24, the integration is performed only in the region of space where *ρ*(*ξ*, *t*) is different from zero. At any given time *t*, Eqs. 23 and 24 are solved iteratively and self-consistently until *C_S_* converges to a specific value. Note that consistently with the criteria stated above, Eq. 23 implies that a negligible error on *ρ*(*ξ*) leads to *ρ*(*ξ*, *t*) = *P*(*ξ*), whereas a large uncertainty on *ρ*(*ξ*) reduces the amount of bias added to the simulation.

In practice, the target distribution *ρ*(*ξ*, *t*) is constantly updated during the simulation, and after a transient period it typically oscillates around an optimal solution. However, a wide confidence band around *ρ*(*ξ*) can result in wide fluctuations of *ρ*(*ξ*, *t*) during the trajectory, potentially compromising the convergence of the method. To avoid such instabilities, a variation of Eq. 23 is used to update *ρ*(*ξ*, *t*) at a slower pace, namely:

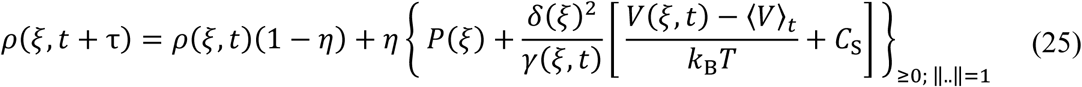

where *τ* is the time range after which *ρ*(*ξ*, *t*) is updated and *0< η < 1* is an update rate (see below).

The parameter *γ*(*ξ*, *t*) in the previous equations is a weight factor of *δ*(*ξ*), leading to an effective noise term 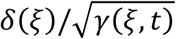. The scale factor *γ*(*ξ*, *t*) is set to attain the largest effective error (i.e. the minimum bias applied to the MD trajectory) that maintains *ρ*(*ξ*, *t*) within the confidence band. This is achieved by imposing the condition |*ρ*(*ξ*, *t*) − *P*(*ξ*)|/*δ*(*ξ*) = 1, from which the following update rule for *γ*(*ξ*, *t*) can be deduced:

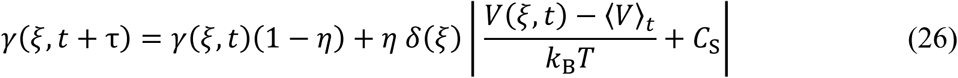

When the uncertainty in *ρ*(*ξ*) is large, update schemes can be also devised for *δ*(*ξ*) and *P*(*ξ*) to further reduce the time oscillations of *ρ*(*ξ*, *t*). For example, if *ρ*(*ξ*, *t*) is within the confidence band, *δ*(*ξ*) can be varied so that it matches approximately the difference between *ρ*(*ξ*, *t*) and *P*(*ξ*). Similarly, *P*(*ξ*) can be updated to get closer to *ρ*(*ξ*, *t*) provided that the latter distribution remains in the confidence band. Further details and specific guidelines for the choice of the simulation parameters are discussed below.

Like in the original version of EBMetaD, the target distribution *ρ*(*ξ*, *t*) is enforced during the MD simulation by adding the biasing potential in Eq. 22 to the energy function. After an equilibration time *t_e_* this potential converges to a well-defined curve, and the resulting stationary distribution *ρ*(*ξ,t >* *t_e_*), calculated over the simulation, approaches the target with the precision dictated by the confidence band. At convergence, the average biasing potential and the calculated probability distribution can be used to deduce the free energy, *G*, as a function of *ξ* (18):

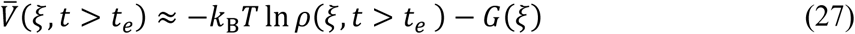

As shown previously (18), the current methodology can simultaneously target multiple probability distributions determined using independent experiments (if these distributions can be assumed to be mutually compatible), simply by summing the corresponding biasing potentials.

Finally, a useful metric to compare different structural interpretations of the experimental data is provided by the reversible work *W* required to produce the biased EBMetaD ensemble. This is related to the total amount of bias added throughout the simulation, at time *t*_s_:

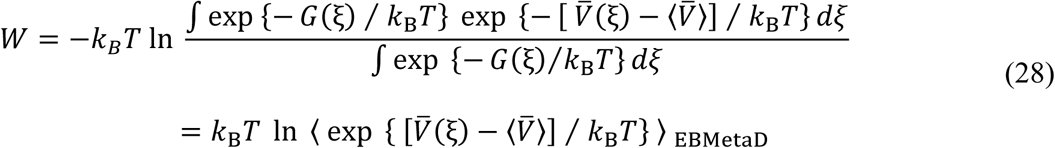

where 〈…〉_EBMetaD_ stands for a time-average over the simulation, again for *t > t_e_*, and:

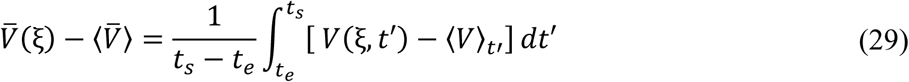

where *t*_s_ is the total simulation time. Note that the value of *W* from Eqs. 28-29 can be derived analogously from the Kullback-Leibler divergence of the probability distributions sampled by EBMetaD and by an unbiased, converged MD simulation, i.e. a measure of the distance between the two ensembles. That is:

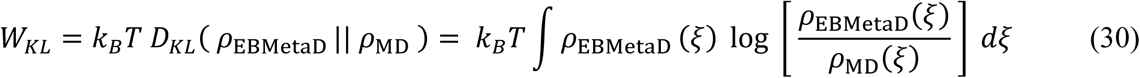

### Ensemble-Biased Metadynamics: Implementation

The extension of EBMetaD described herein can be freely used with NAMD 2.12 (28) and LAMPS (37), specifically through the “colvars” module (38). This implementation follows the formalism introduced above. As mentioned, the convergence of this technique is related to the fluctuations of the target probability density *ρ*(*ξ*, *t*) during the trajectory. The time variability of this distribution depends on the value of parameters *w, σ_G_* and *τ* in Eq. 22, on the update rate of *ρ*(*ξ*, *t*) and on the width of the confidence band (*P*(*ξ* ± *δ*(*ξ*). To optimize the performance of EBMetaD, *ρ*(*ξ*) and *δ*(*ξ*) may be updated on time according to the following criteria:

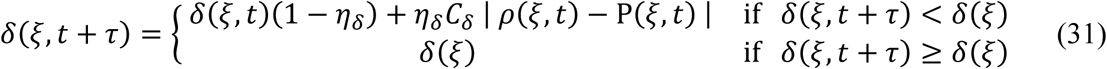

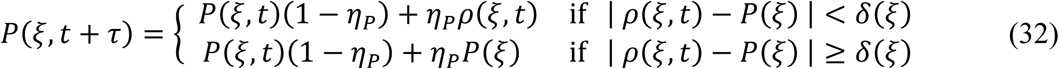

In these equations *C_δ_* >1, and was set to 1.5 for all the simulations. The parameters η*_δ_* and *η_δ_* are update rates (0< *η_δ_, η_P_* <1) that must be selected as a fraction of *η* in Eqs. 31-32. The latter term is also selected as a fraction (0≤*C_η_* ≤ 1) of the biasing potential update rate (18):

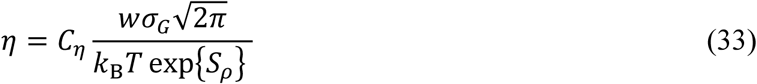

The choice of *η, η_δ_* and *η_P_* in Eqs. 31-33 relates to the time required to reach equilibration; the duration of this equilibration stage is on the order of *τ* divided by the corresponding rate parameter. To accelerate convergence, the rate parameters can be selected larger in the first part of the simulation and then gradually reduced. To assess whether the latter parameters have been set reasonably, it is useful to monitor the time fluctuations of the constant *C*_S_ and of *ρ*(*ξ*, *t*) (Eqs. 23-24). Large oscillations in *C*_S_, associated with intermittent values of *ρ*(*ξ*, *t*) that become zero, are an indication of poor convergence, implying that the value of the rate parameters must be reduced.

For the butyramide simulation, the variable biased by EBMetaD is the dihedral angle defined by atoms N, C, C_α_ and C_β_. Gaussians of height *w* = 0.025 kcal/mol and width *σ_G_* = 5° were added every 2 ps and were scaled by the target distribution according to Eq. 22. The rate parameters were set as *C_η_* = 1, = *η_δ_* = *η_P_* = *η/*10. During the initial equilibration stage, lasting 30 ns, the Gaussian height and the rate parameters were gradually reduced to *w* = 0.01 kcal/mol, *η_δ_ =η*/10 and *η_P_* = *η*/40.

In the simulation of T4L the variables biased by EBMetaD are the distances between the centers-of-mass of the nitroxide groups in the spin-labels. Gaussians of width *σ_G_* = 0.5 Å were added every 2 ps. The parameter *w* was initially set to 0.05 kcal/mol and gradually reduced to 0.01 kcal/mol during equilibration (first 100 ns of simulation). The sampling of the spin labels distance was restricted using flat-bottom potentials in the ranges [21.8 Å, 38.6 Å], [37.2 Å, 46.3 Å], [12.3 Å, 48.8 Å] for spin-label pairs 62-109, 62-134 and 109-134 respectively. To avoid the onset of systematic errors at the boundaries of these intervals, the Gaussians added to the biasing potential were reflected beyond the boundaries (39), which translates into a flat biasing potential at the ends of those intervals. Accordingly, the biasing forces were set to zero outside the boundaries. In the EBMetaD simulations including the confidence band, the rate parameters were set according to *C_η_* = 0.25, = *η_δ_* = *η_P_* = *η/*10 and then in the production run they were scaled down to *η_δ_* = *η*/10 and *η_P_* = *η*/40.

## RESULTS

### Influence of Noise Level on the Confidence Band for a *P*(*R*)

We first evaluate using simulated DEER signals how the noise level of the data is reflected in the estimated uncertainties of the fit parameters and the confidence band for the distance distribution. **Fig. 1** shows results obtained from fitting a simulated signal at two different noise levels using DD (https://lab.vanderbilt.edu/hustedt-lab/software). Consistent with the fact that they were simulated for an unimodal distance distribution, the *n* = 1 model gives lower *BIC* values for both data sets and is thus favored (**Table 1**). The best-fit P(R) for the low noise example agrees very well with the true distribution, while the best-fit *P*(*R*) for the high noise case is shifted from the true distribution due to the higher variance in the fit parameters.

**Table 1.** Model selection for fits in Fig. 1.

| *noise* | *n* | $\chi^2_v$ | $\Delta BIC$ |
| --- | --- | --- | --- |
| 0.005 | 1 | 0.984 | 0.0 |
|  | 2 | 0.982 | 13.3 |
| 0.050 | 1 | 1.054 | 0.0 |
|  | 2 | 1.0581 | 15.5 |

**Figure 1.**
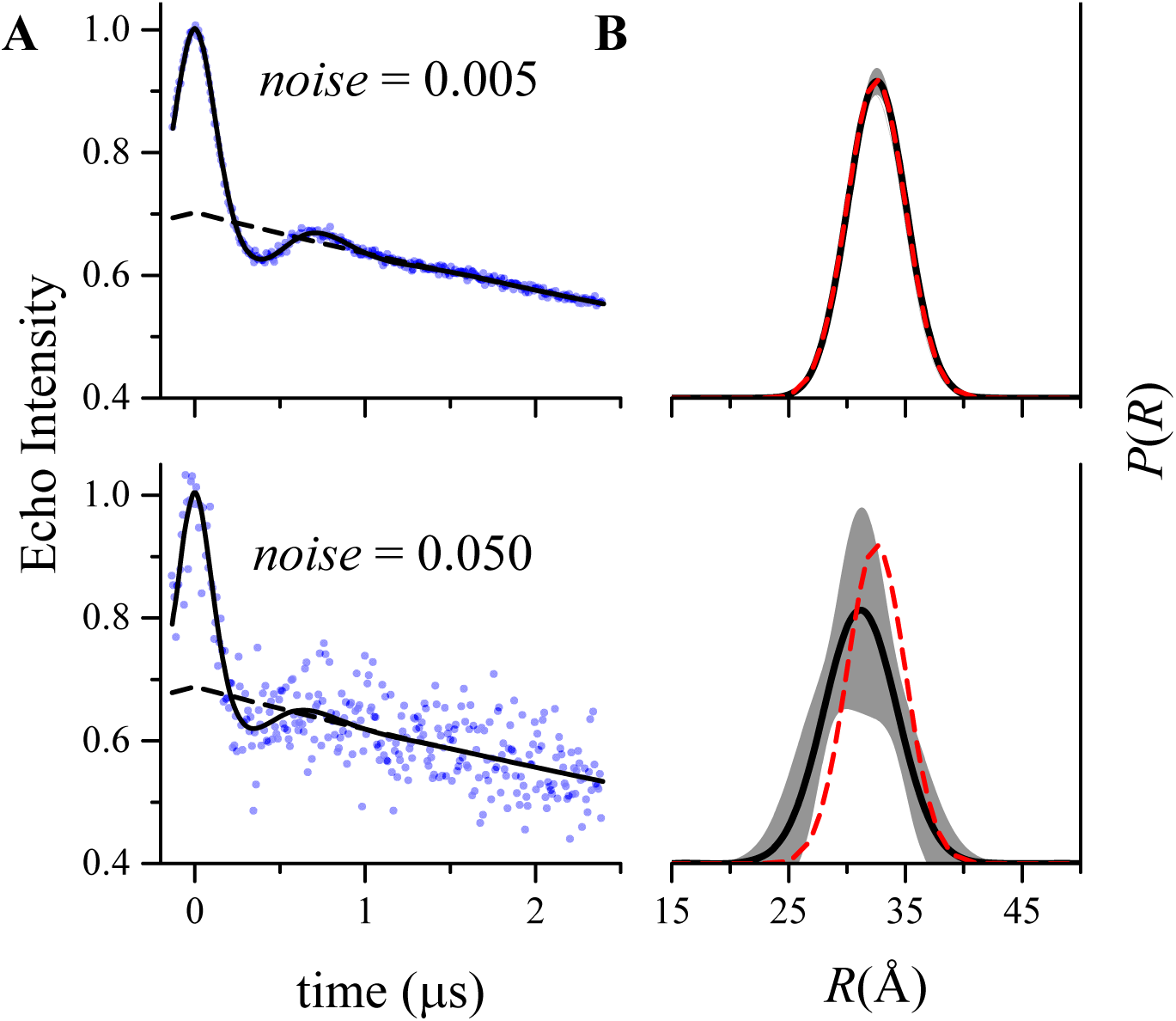
Fits to simulated DEER data generated using a single Gaussian to model *P*(*R*) (*r*_0_ = 32.5 Å and *σ_r_* = 2.5 Å). Data were simulated for *t* = −128 ns to +2400 ns with a time increment of 8 ns. Normally-distributed random numbers with standard deviation of either 0.005 or 0.050 were added as noise. A) The simulated data (blue dots), the fits (solid black lines), and the best-fit background factor (dashed black lines). B) The best fit *P*(*R*) (solid black lines), the confidence band (2σ, shaded grey regions), and the true *P*(*R*) (dashed red lines) used to generate the simulated signal. Only a portion of the full range (0 – 100 Å) of *R* is shown. The values of each of the fit parameters and their uncertainties are given in **Table 2**.

The best-fit parameters from these fits are given in **Table 2** along with the parameter uncertainties estimated from the covariance matrix (Eqs. 15-17) and the upper and lower parameter bounds estimated from confidence-interval calculations. Example confidence intervals for the parameters r_0_ and *σ_r_* are shown in **Fig. 2A**. The values reported for the upper and lower parameter limits are measured where each 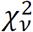 curve intersects the green dashed line corresponding to the 2*σ* confidence level.

**Table 2.**
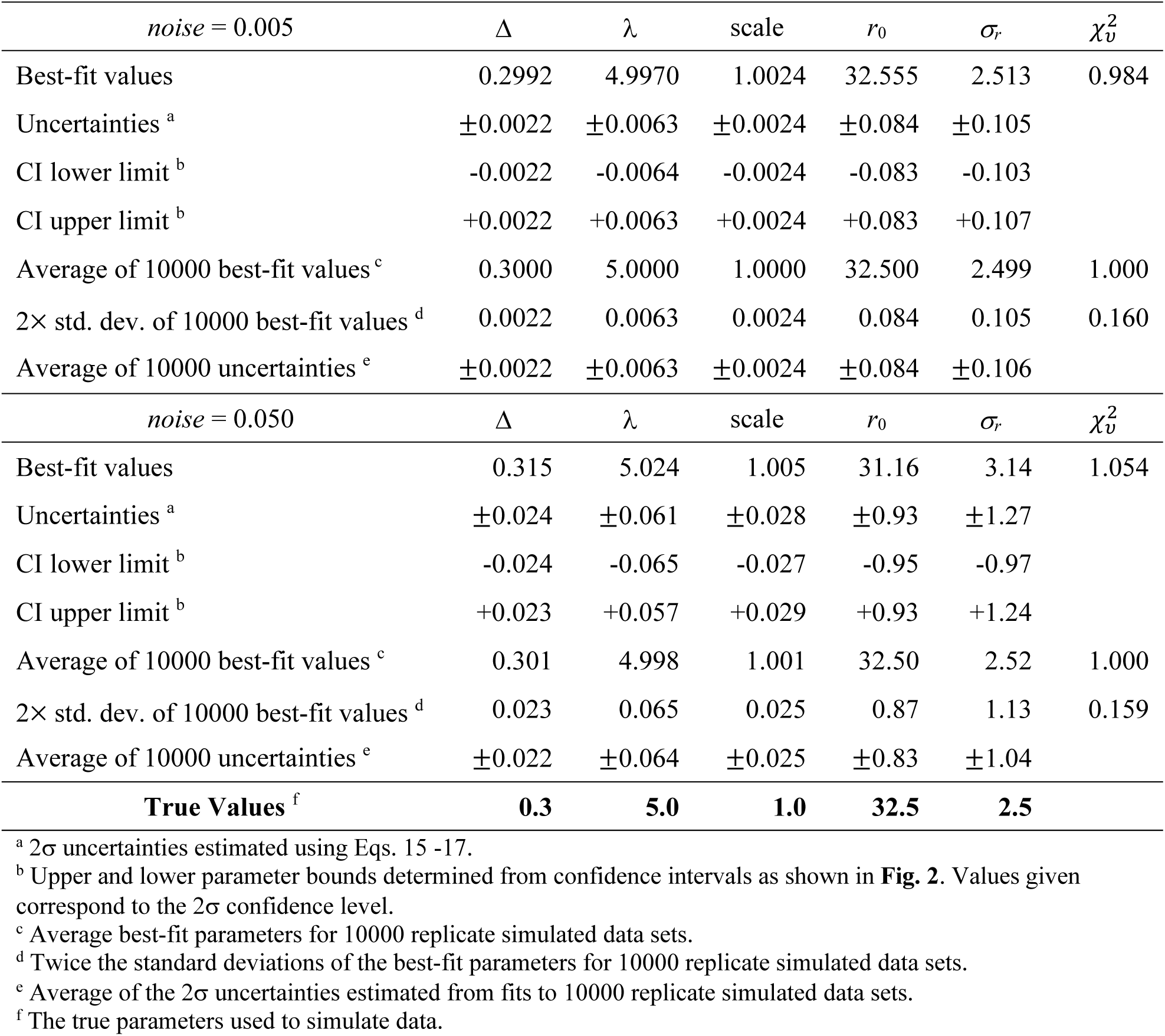
Best-fit parameters for the simulated DEER signals in Fig. 1.

| *noise* = 0.005 | $\Delta$ | $\lambda$ | scale | $r_0$ | $\sigma_r$ | $\chi^2_v$ |
| --- | --- | --- | --- | --- | --- | --- |
| Best-fit values | 0.2992 | 4.9970 | 1.0024 | 32.555 | 2.513 | 0.984 |
| Uncertainties <sup>a</sup> | $\pm 0.0022$ | $\pm 0.0063$ | $\pm 0.0024$ | $\pm 0.084$ | $\pm 0.105$ | |
| CI lower limit <sup>b</sup> | -0.0022 | -0.0064 | -0.0024 | -0.083 | -0.103 |  |
| CI upper limit <sup>b</sup> | +0.0022 | +0.0063 | +0.0024 | +0.083 | +0.107 |  |
| Average of 10000 best-fit values <sup>c</sup> | 0.3000 | 5.0000 | 1.0000 | 32.500 | 2.499 | 1.000 |
| 2× std. dev. of 10000 best-fit values <sup>d</sup> | 0.0022 | 0.0063 | 0.0024 | 0.084 | 0.105 | 0.160 |
| Average of 10000 uncertainties <sup>e</sup> | $\pm 0.0022$ | $\pm 0.0063$ | $\pm 0.0024$ | $\pm 0.084$ | $\pm 0.106$ | |
| *noise* = 0.050 | $\Delta$ | $\lambda$ | scale | $r_0$ | $\sigma_r$ | $\chi^2_v$ |
| Best-fit values | 0.315 | 5.024 | 1.005 | 31.16 | 3.14 | 1.054 |
| Uncertainties <sup>a</sup> | $\pm 0.024$ | $\pm 0.061$ | $\pm 0.028$ | $\pm 0.93$ | $\pm 1.27$ | |
| CI lower limit <sup>b</sup> | -0.024 | -0.065 | -0.027 | -0.95 | -0.97 |  |
| CI upper limit <sup>b</sup> | +0.023 | +0.057 | +0.029 | +0.93 | +1.24 |  |
| Average of 10000 best-fit values <sup>c</sup> | 0.301 | 4.998 | 1.001 | 32.50 | 2.52 | 1.000 |
| 2× std. dev. of 10000 best-fit values <sup>d</sup> | 0.023 | 0.065 | 0.025 | 0.87 | 1.13 | 0.159 |
| Average of 10000 uncertainties <sup>e</sup> | $\pm 0.022$ | $\pm 0.064$ | $\pm 0.025$ | $\pm 0.83$ | $\pm 1.04$ | |
| **True Values <sup>f</sup>** | **0.3** | **5.0** | **1.0** | **32.5** | **2.5** |  |

<sup>a</sup> 2σ uncertainties estimated using Eqs. 15 -17.
<sup>b</sup> Upper and lower parameter bounds determined from confidence intervals as shown in **Fig. 2**. Values given correspond to the 2σ confidence level.
<sup>c</sup> Average best-fit parameters for 10000 replicate simulated data sets.
<sup>d</sup> Twice the standard deviations of the best-fit parameters for 10000 replicate simulated data sets.
<sup>e</sup> Average of the 2σ uncertainties estimated from fits to 10000 replicate simulated data sets.
<sup>f</sup> The true parameters used to simulate data.

**Figure 2.**
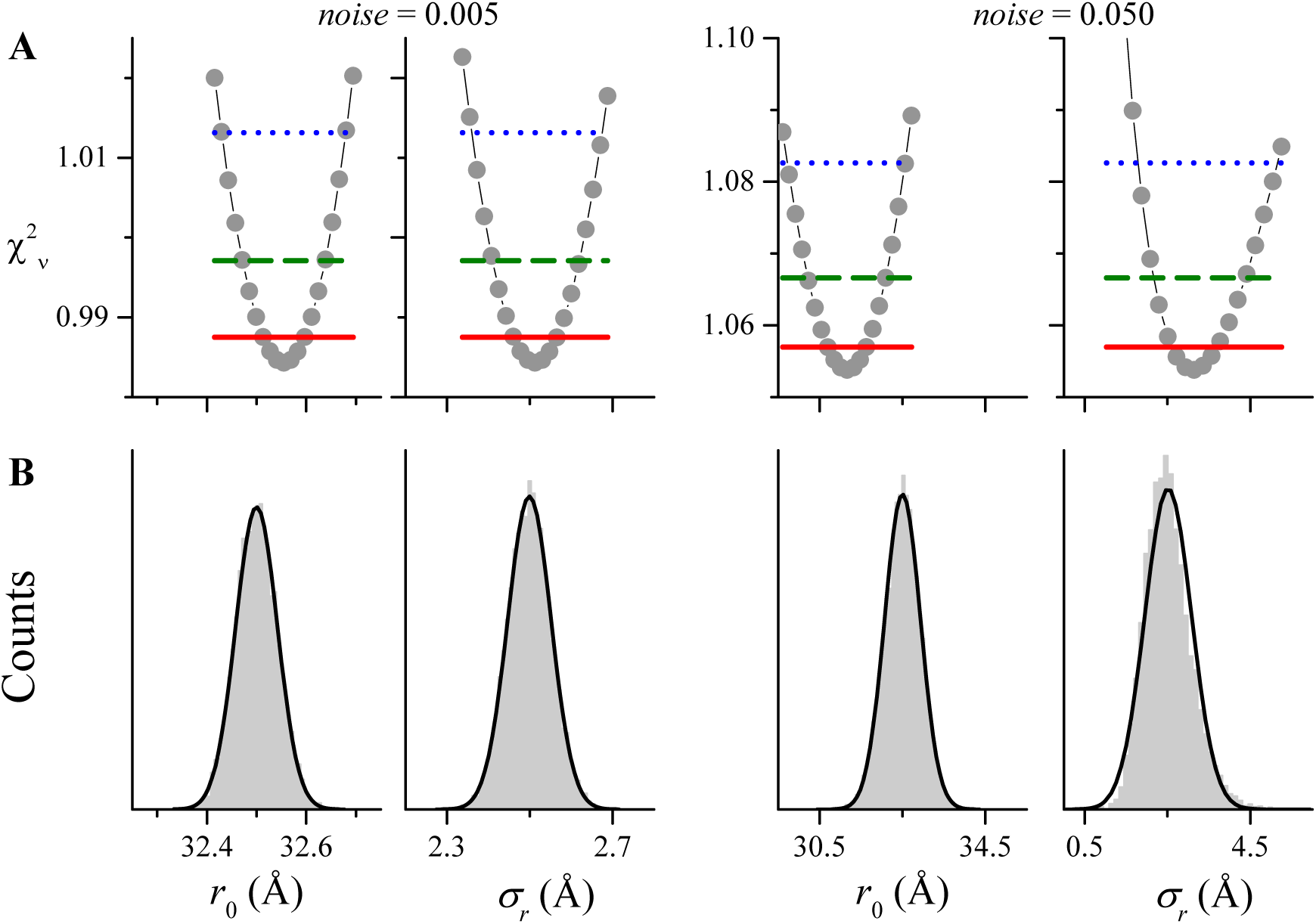
Comparison of 1-dimensional confidence intervals (A) for *r*_0_ and *σ_r_* from the fits in Fig. 1 to histograms (B) from fitting 10000 replicate data sets. The 4 panels on the left were obtained for the lower noise level (0.005); the 4 on the right were obtained for the higher noise level (0.05). A) Grey dots obtained by fixing the parameter *r*_0_ or *σ_r_* to a series of values and allowing the other four fit parameters to vary to minimize 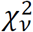. The horizontal lines give the 1σ (solid red), 2σ (dashed green), and 3σ (dotted blue) confidence levels. Lower and upper bounds on the parameters at a particular confidence level are determined by where the 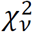 curve intersects the appropriate horizontal line. B) Histograms of 10000 parameter values obtained from repetitive fits to data similar to that in **Fig. 1**. The solid black lines are normal (Gaussian) distributions calculated for the mean and standard deviation of the distribution of parameter values (see **Table 2**).

To ascertain the validity of these parameter uncertainties and confidence bands, we analyzed fits for 10000 replicate signals generated using a Monte Carlo procedure from the same model and with the same level of added random noise. Examining all of these results, three important conclusions can be drawn. First, the distribution of parameter values obtained from 10000 replicate fits are typically well-described by a Gaussian distribution (**Fig. 2B**) and that the parameter uncertainties estimated from a single fit (at the 2*σ* confidence level) match (twice) the value of the standard deviation of these parameter distributions (**Table 2**) as expected. At the highest noise level, the best-fit parameters for a fit to a single data set are shifted from the true values due to the increase in the parameter variance and both the confidence interval from the singe fit and the histogram from 10000 fits for the *σ_r_* parameter are slightly distorted by the zero lower bound on the parameter (**Fig. 2**, far right). Second, as long as the errors are Gaussian, the uncertainties estimated from the covariance matrix match the results of the more rigorous confidence interval calculations. Finally, the parameter uncertainties increase linearly as the noise level increases.

Confidence bands, *P*(*R*) *±* 2*δ*(*R*) (Eqs. 18-21), for the best-fit distance distributions are shown in **Fig. 1B** as grey shaded regions, and the *δ*(*R*) themselves are plotted in **Fig. 3** (red dotted lines). **Fig. 3** also includes the standard deviation of the *P*(*R*) obtained from fitting 10000 replicate data sets (solid grey lines) and the average value of *δ*(*R*) from these fits (dashed black lines). The results for the lower noise level (**Fig. 3**, upper) show that the *δ*(*R*) obtained from fitting a single data set overlays the standard deviation in *P*(*R*) that would be obtained from fitting a large number of replicate data sets. The results at the higher noise level (**Fig. 3**, lower) show that the *δ*(*R*) from a single fit gives a reasonable order-of magnitude estimate of this standard deviation which depends linearly on the noise level.

**Figure 3.**
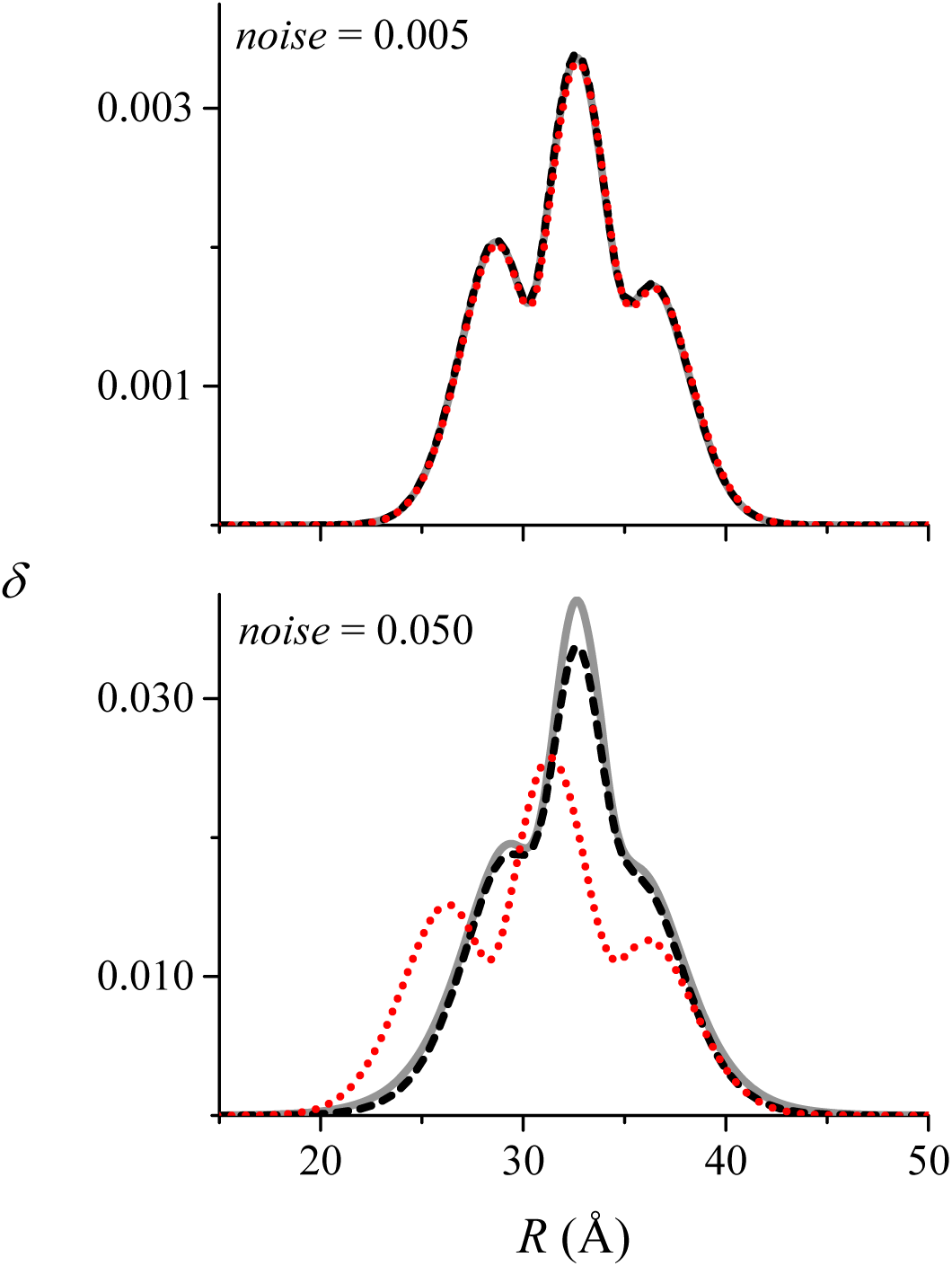
Comparison of the *δ*(*R*) (red dotted line) calculated (Eqs. 18-21) for the fits in Fig. 1 to the average *δ*(*R*) (black dashed line) and the standard deviation (solid grey line) of all of the *P*(*R*) obtained from fitting 10000 replicate data sets. The results in the upper panel were obtained for the lower noise level (0.005) and the results in the lower panel for the higher noise level (0.05). Only a portion of the full range (0 – 100 Å) of *R* is shown.

In summary, the results shown in Figs. 1-3 demonstrate that the parameter uncertainties and the confidence bands for *P*(*R*) both properly account for the noise in the data and give reasonable estimates of the distributions that would be obtained from fitting multiple replicate data sets.

### Background Correction Uncertainty

In addition to random noise, other factors can influence the magnitude of parameter uncertainties and confidence band for P(R). In particular, the maximum observed dipolar evolution time determines to what degree the background factor can be resolved at the tail of the full DEER signal. In **Fig. 4**, a “stress test” is performed using DD to fit simulated DEER signal with an extremely short dipolar evolution time generated using the same model parameters and noise level as that in **Fig. 1** (upper). The simulated data are well-fit using a single Gaussian to model P(R) while a two-component model gives a larger *BIC* value (see **Supplemental Table S1**).

**Figure 4.**
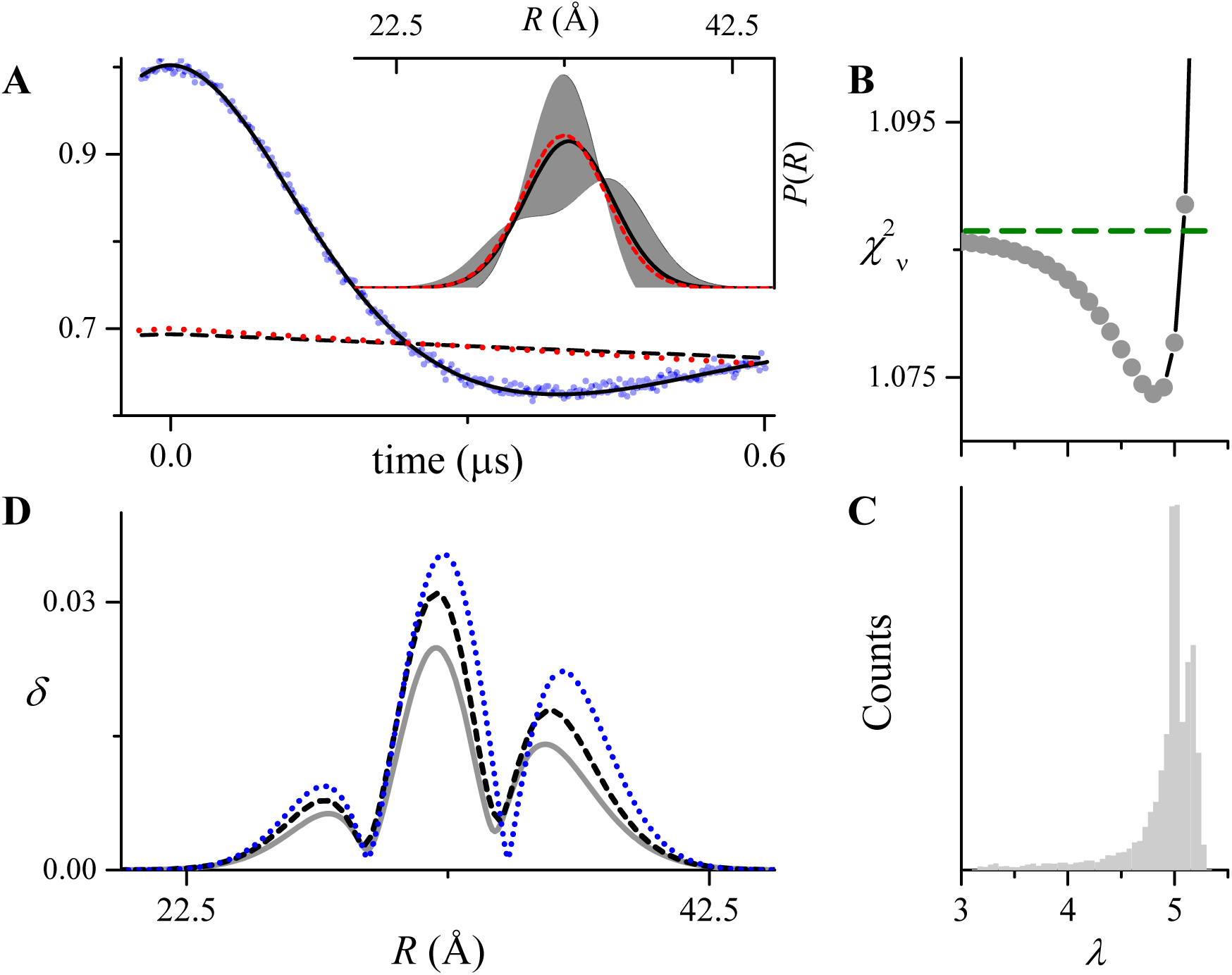
Fit to a simulated DEER data generated using a single Gaussian to model *P*(*R*) (*r*_0_ = 32.5 Å and *σ_r_* = 2.5 Å) and a short dipolar evolution time. Data was simulated for *t* = −32 ns to +600 ns with a time increment of 2 ns. Other parameters are given in **Supplemental Table S2**. Normally-distributed random numbers with standard deviation of 0.005 were added as noise. A) The simulated data (blue dots), the fit (solid black line), the best-fit background factor (dashed black line), and the true background (dotted red line). The inset shows the best-fit *P*(*R*) (solid black lines), the confidence band (2σ, shaded grey regions), and the true *P*(*R*) (dashed red lines) for the simulated data. B) One-dimensional confidence interval (grey dots) obtained by fixing *λ* to a series of values and allowing the other four fit parameters to vary to minimize 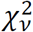. The green dashed horizontal line gives the 2σ confidence level. C) Histograms of 10000 *λ* values obtained from repetitive fits to replicate data similar to that in panel A. D) The *δ*(*R*) (blue dotted line) calculated for the fit A compared to the average *δ*(*R*) (black dashed line) and the standard deviation (solid grey line) of all of the *P*(*R*) obtained from fitting 10000 replicate data sets. Only a portion of the full range (0 – 100 Å) of *R* is shown.

Despite the fact that the background is not resolved for this simulated signal, DD is able to determine a reasonable estimate of the background correction as can be seen by comparing the best-fit background (**Fig. 4A**, dashed black line) with the true background factor (dotted red line) or by comparing the best-fit *λ* and Δ parameters to the true values (see **Supplemental Table S2**). Nonetheless, there is considerable uncertainty in the parameter *λ* as determined by either the covariance matrix (4.83 ± 0.88) or the 1-dimensional confidence interval (**Fig. 4B**). This confidence interval for *λ* and those for most of the other fit parameters (not shown) strongly deviate from the parabolic shapes seen in **Fig. 2**. Accordingly, analysis of the 10000 replicate signals reveals a broad range of *λ* values (**Fig. 4C**), which in turn leads to large variations in the other fit parameters (**Supplemental Fig. S1**). Most of these histograms strongly deviate from the Gaussian shapes seen in **Fig. 2**. The lack of precision in determining the background correction leads to a confidence band for *P*(*R*) that is dramatically larger than that obtained for data collected for a longer dipolar evolution time (*cf*. **Fig. 1** upper panel). However, the calculated *δ*(*R*) (dotted blue line) give a reasonable estimate of the standard deviation in *P*(*R*) that would be obtained from fitting a large number of replicate data sets (solid grey line). Finally, even under the extreme conditions presented by the simulated data in **Fig. 4A**, the best-fit *P*(*R*) is very close to the true *P*(*R*) demonstrating that the distance distribution and the background factor can be simultaneously estimated using our approach.

### Multimodal Gaussian Distribution

Of critical importance is the performance of the proposed analysis method for DEER signals originating from multi-modal distance distributions as is typical for systems of biological interest. Fits to simulated DEER signals calculated for two different trimodal distance distributions are shown in **Fig. 5A**. As expected for both data sets the optimal model based on *BIC* values is a sum of 3 Gaussians (*n* = 3, **Supplemental Table S3**). The parameter uncertainties estimated from the covariance matrix are in excellent agreement with the results from 1-dimensional confidence-interval calculations and with the standard deviations of the parameter values resulting from fitting 10000 replicate signals (**Supplemental Table S4**). Likewise, the *δ*(*R*) used to calculate the confidence band for *P*(*R*) for each fit closely agrees with the standard deviation of *P*(*R*) obtained from fitting 10000 replicates (**Fig. 5B**). For both simulated signals, neither the parameter uncertainties nor the widths of the confidence bands depend on the *r*_0_ values of the individual components in a straightforward way. For the data in the upper panel of **Fig. 5A** for which the true *σ_r_* values of the three Gaussians are equal, the parameter uncertainties and width of the confidence band are roughly equal for the three components. For the data in the lower panel of **Fig. 5A** the parameter uncertainties for r_0_ and *σ_r_* increase as the value of *σ_r_* increases for the three components. However, the width of the confidence band decreases as *σ_r_* increases.

**Figure 5.**
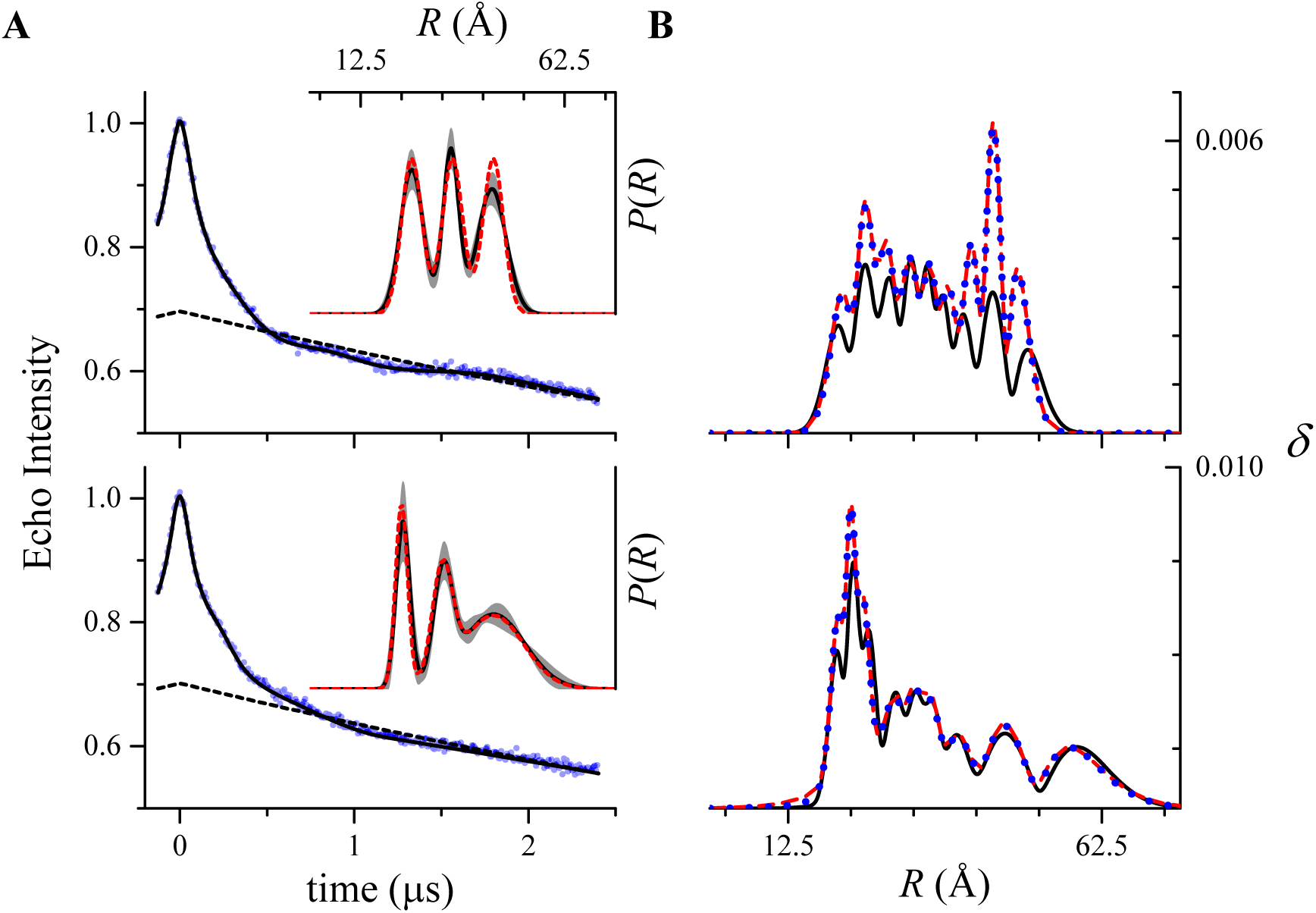
Fits to two simulated DEER signals generated using three Gaussians to model *P*(*R*). Data were simulated for *t* = −128 ns to +2400 ns with a time increment of 8 ns.Other parameters are given in **Supplemental Table S4**. Normally-distributed random numbers with standard deviation of 0.005 were added as noise. A) The simulated data (blue dots), the fits (solid black lines), and the best-fit background factor (dashed black lines). The insets show the best fit *P*(*R*) (solid black lines), the confidence bands (2σ, shaded grey regions), and the true *P*(*R*) (dashed red lines) for the simulated data. B) Comparison of the *δ*(*R*) (solid black lines) calculated for the fits in A to the average *δ*(*R*) (dotted blue lines) and the standard deviation (dashed red lines) of all of the *P*(*R*) obtained from fitting 10000 replicate data sets. Only a portion of the full range (0 – 100 Å) of *R* is shown.

To further clarify how the uncertainty in a component of *P*(*R*) varies as its average distance value increases, additional calculations for a unimodal simulated signal were performed as summarized in **Supplemental Table S5**. For *r*_0_ up to 45 Å the uncertainties in *r*_0_ and *σ_r_* do not vary significantly, while for *r*_0_ = 55 Å and beyond, for which a full modulation period is not completed within the dipolar evolution time of 2400 ns, the uncertainties in both *r*_0_ and *σ_r_* increase dramatically.

In summary, the results in **Fig. 5** together with Supplemental Tables S3 and S4 demonstrate that all of the new methods presented here perform as expected when DEER data is derived from complex, multimodal distributions.

### Comparison to DeerAnalysis

For comparison with the TR method, fits to the simulated signals in **Fig. 1** obtained using DeerAnalysis2016 are shown in **Supplemental Figs. S2 and S3**. At the lower noise level, the two approaches give similar results and similar estimates of the uncertainty in *P*(*R*). At the higher noise level, the DD estimate is considerably larger. This may be due, at least in part, to the fact that, as is commonly done, only the background starting time option was used here in the validation tool of DeerAnalysis2016. For the simulated signals in Fig. 4, it is difficult to find a convincing *a priori* background correction, given the short dipolar evolution time of the data, and therefore this signal cannot be interpreted using DeerAnalysis2016. For the multi-component simulated signals in **Fig. 5**, fits obtained with DeerAnalysis2016 are shown in **Supplemental Figs. S4 and S5**. For the signal generated from a *P*(*R*) calculated as the sum of three Gaussians of equal width, both DD and DeerAnalysis2016 give similar results.

For the data generated from a *P*(*R*) using 3 Gaussians of varying widths, DeerAnalysis2016 adds a fourth component at *r*_0_ ≈ 40 Å, apparently to account for the component with the broadest width (**Supplemental Fig. S5**). This result is due to the fact that TR tends to produce multicomponent distributions with equal component widths. It is important to note that the confidence bands for *P*(*R*) obtained from DD (**Supplemental Fig. S4**F **and S5F**) do not strictly follow the color coding scheme for reliability in DeerAnalysis2016 (**Supplemental Fig. S4D and S5D**).

### Evaluation of generalized EBMetaD for a small-molecule system

Having established the validity of our signal analysis algorithm, we evaluated the performance and accuracy of the generalized EBMetaD method. For this purpose, we first considered a simple system, namely a butyramide molecule in water. The conformational descriptor considered in this evaluation is the dihedral angle (Ψ) around the C_α_-C bond i.e. *ξ* = Ψ in Eq. 22 (**Fig. 6**). We first calculated a 400-ns trajectory using a standard MD simulation, so as to obtain a well-converged probability distribution along Ψ. This distribution is symmetric around 0° with peaks at Ψ ≈ ±70° (**Fig. 6**). We then designed an artificial target distribution for the EBMetaD method, substantially different from that obtained above, as well as several confidence bands of increasing width around this target distribution, such that the widest of these bands encompass the unbiased MD distribution, partially or fully (**Fig. 6**). Using each of these confidence bands as input, we then calculated a 320-ns trajectory with EBMetaD using the same simulation parameters employed for the conventional MD trajectory.

**Figure 6.**
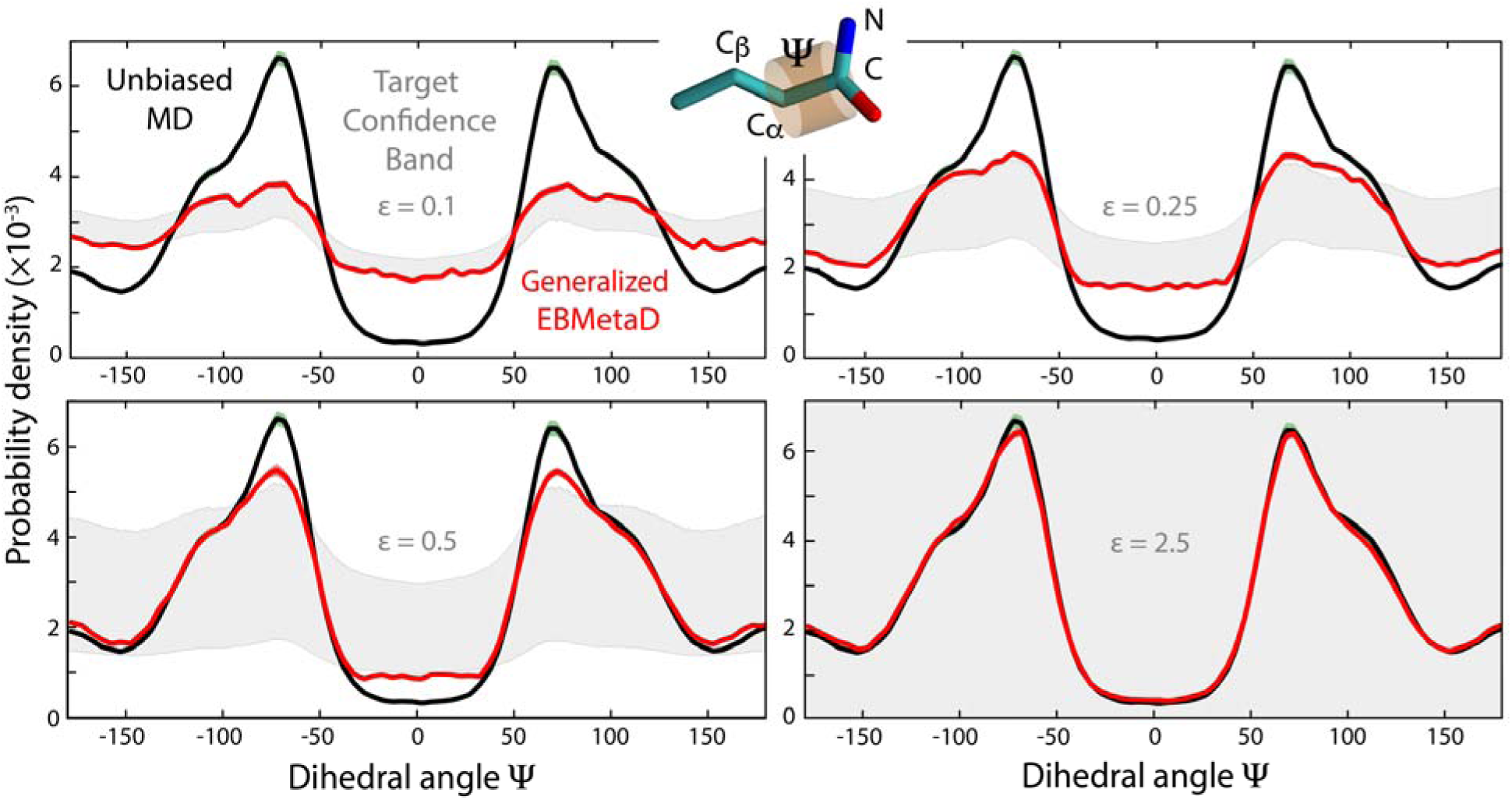
Evaluation of the generalized EBMetaD method for butyramide in water. The torsional angle Ψ defined by atoms C_β_, C_α_, C and N is considered. The probability distribution obtained from conventional, unbiased MD simulation (*black*), *ρ*_MD_(Ψ), is compared with those obtained using EBMetaD simulations (*red*), *ρ*_EBMetaD_(Ψ), in four independent calculations that target a different, hypothetical distribution and its uncertainty band (*grey*), constructed as *P*(Ψ) ± *εP*(Ψ), where *ε* = 0.1, 0.25, 0.5, 2.5. Note that *ρ*_MD_(Ψ) is partially encompassed by the target confidence band at *ε* = 0.5, and fully encompassed at *ε* = 2.5.

The distributions along Ψ obtained after equilibration illustrate the performance of the proposed method. The EBMetaD distributions draw along the edges of the confidence band, so as to fulfill the target data while remaining as close as possible to the unbiased distribution (**Fig. 6**). Accordingly, when the confidence band is wide enough to encompass the unbiased probability density, the EBMetaD approach does not bias the sampling, and produces the same distribution as standard MD. By contrast, when the confidence band is narrow, the EBMetaD distribution deviates from the unbiased distribution as needed.

Analysis of the converged biasing potential developed by the EBMetaD algorithm for each of the input confidence bands, using Eq. 29, explains these results (**Fig. 7A**). It is apparent that the magnitude of the bias added to the simulation is not uniform along Ψ, but depends on the difference between the target and unbiased distribution. The overall bias also decreases as the uncertainty of the target distribution is greater; when the input confidence band encompasses the unbiased distribution from conventional MD, the EBMetaD potential is nearly flat. i.e. no conformational bias is applied (**Fig. 7A**). This result, consistent with the minimum-information condition, can be further quantified by calculating the reversible work performed to enforce the EBMetaD biasing potential, using Eq. 28 (**Fig. 7B**). Consistent with the data in **Fig. 7A**, the value of the work diminishes as the confidence band widens, becoming negligible when the unbiased MD distribution is within the uncertainty.

**Figure 7.**
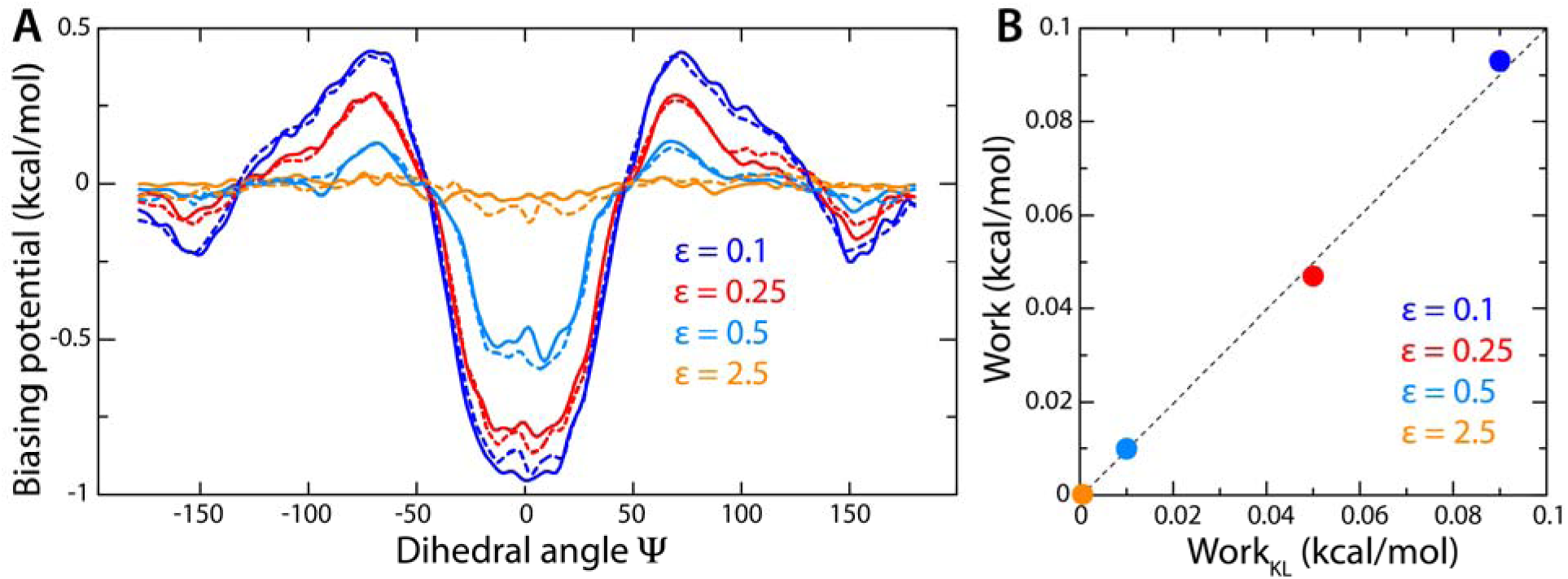
Evaluation of the generalized EBMetaD method for butyramide in water. (**A**) EBMetaD biasing potential as function of Ψ, for each of the calculations shown in **Fig. 6**, i.e. with increasing confidence-band widths (denoted by the scaling factor *ε*). The biasing potential is calculated both Eq. 29, with *t_e_* = 30 ns and *t* = 320 ns (*solid curves*), and compared with a calculation using Eq. 27 (dashed lines), in which *G*(Ψ) = −*k_B_Tlnρ*_MD_(Ψ), where *ρ*_MD_(Ψ) is the unbiased probability distribution obtained from an unbiased MD trajectory (**Fig. 6**). (**B**) Reversible work required to enforce the EBMetaD biasing potential, for each of the four simulations abovementioned. The work values are derived from the Kullback-Leibler divergence of *ρ*_MD_(Ψ) and *ρ*_EBMetaD_(Ψ) (**Fig. 6**), using Eq. 30, and compared with calculations based on the biasing potentials in panel (A), using Eq. 28. The error bars in the work values, based on from a 5-block analysis, range from 10^−3^ to 10^−4^ kcal/mol.

Like with any enhanced-sampling simulation method, it is important for EBMetaD to preserve the inherent thermodynamics of the molecular system. To evaluate whether this is the case, the biasing potentials derived with Eq. 29 were compared with calculations based on Eq. 27, i.e. from the free-energy function = *G*(Ψ) = −*k*_B_*T* ln *ρ*_MD_(Ψ), which for this simple system can be derived from an unbiased trajectory (**Fig. 7A**). Similarly, the work values calculated using Eq. 28 were contrasted with those deduced from the Kullback-Leibler divergence (Eq. 30) of the unbiased and biased distributions (**Fig. 7B**). Both evaluations demonstrate the proposed methodology is robust qualitatively and quantitatively.

In summary, this application demonstrates that EBMetaD constructs a conformational ensemble compatible with the confidence band of a target probability distribution and with the underlying free-energy landscape of the system, and that it does so by applying the minimum bias required. This application also shows that the EBMetaD work is an accurate descriptor of the degree to which an unbiased conformational ensemble might be compatible with a given set of target data. We posit that these features make the EBMetaD method an ideal tool to formulate well-founded molecular interpretations for a range of experimental information e.g. DEER data.

### Model-based fitting and EBMetaD simulations for T4L DEER data

After demonstrating the validity of the extended EBMetaD method on a simple system, we applied this approach to spin-labeled T4L in explicit water. Following our previous work (18), we considered three doubly-labeled T4L mutants with labels introduced at positions 62, 109 and 134. Experimental DEER data along with the corresponding fits obtained using DD are shown in **Fig. 8**. Based on BIC values, a 3-Gaussian model was found to be optimal for T4L 62/109 while a 2-Gaussian model was better suited for both T4L 62/134 and T4L 109/134 (**Table 3**). The best-fit parameters are given in **Supplemental Table S6**. Note that the confidence bands for pairs T4L 62/109 and T4L 109/134 are relatively narrow over the entire distance range while that for T4L 62/134 is very broad for the component at the longest distance. These data sets, therefore, constitute a non-trivial test case of EBMetaD methodology.

**Table 3.** Model selection for T4L data sets.

| T4L mutant | $n$ | $\chi^2_v$ | $\Delta BIC$ |
| --- | --- | --- | --- |
| 62/109 | 1 | 3.741 | 377.0 |
| 62/109 | 2 | 0.713 | 4.1 |
| 62/109 | 3 | 0.662 | 0.0 |
| 62/109 | 4 | 0.654 | 10.5 |
| 62/134 | 1 | 1.395 | 9.4 |
| 62/134 | 2 | 1.220 | 0.0 |
| 62/134 | 3 | 1.243 | 15.1 |
| 109/134 | 1 | 1.400 | 25.8 |
| 109/134 | 2 | 1.049 | 0.0 |
| 109/134 | 3 | 0.977 | 2.1 |

**Figure 8.**
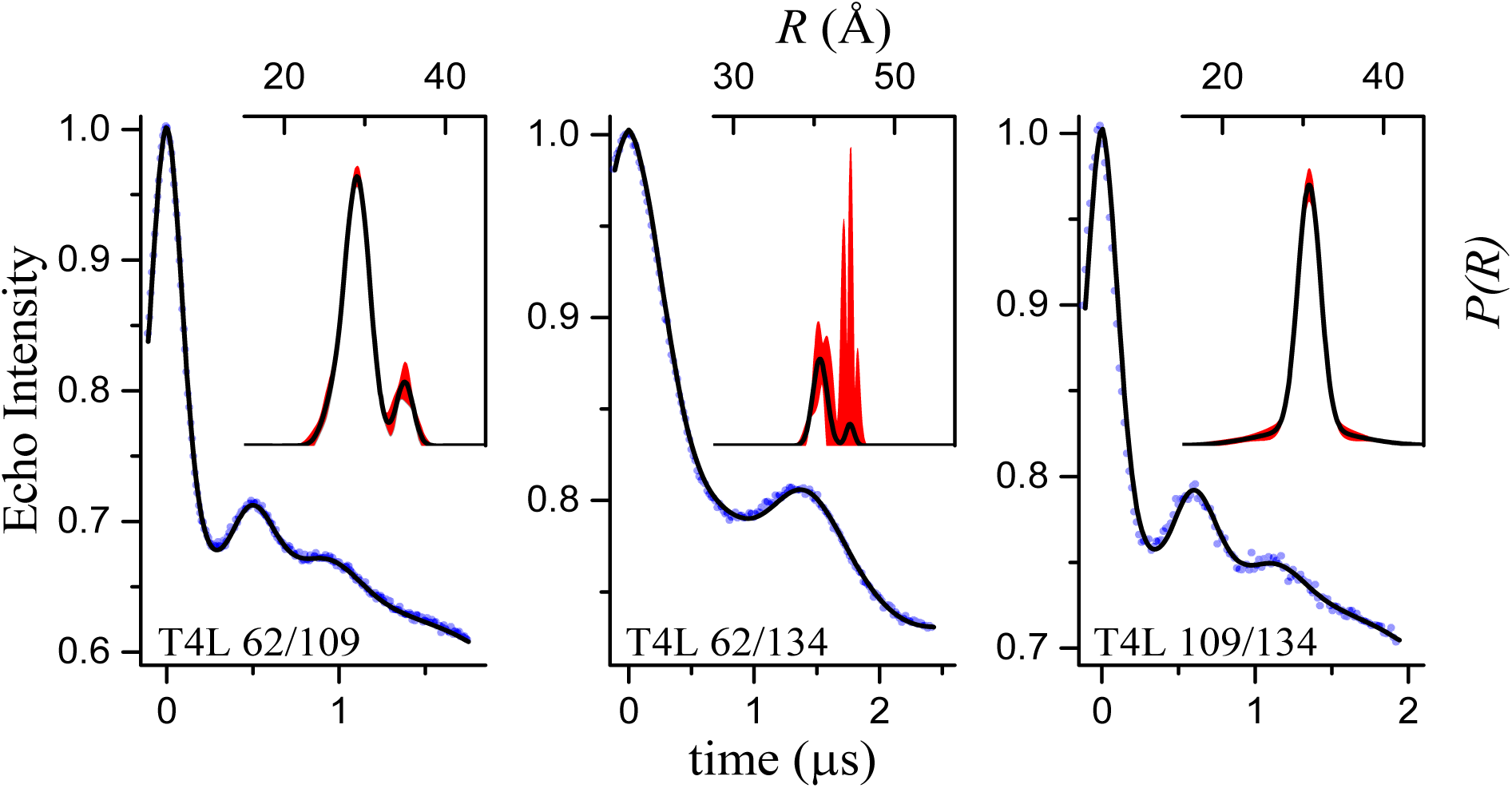
DEER signals and associated probability distributions for three spin-label pairs in T4-lysozyme. The DEER data are shown as blue dots and the fits as solid black lines. The insets show the best fit *P*(*R*) (solid black lines), along with the newly obtained confidence bands (2σ, shaded red regions). Best-fit parameters are given in **Supplemental Table S6**.

To model the DEER data for T4L using EBMetaD, we considered the distance between the centers-of-mass of the nitroxide groups as the reaction coordinate (*ξ* = *R*). For simplicity, we enforced the three experimental distance distributions simultaneously, even though the DEER signals were measured for one pair of spin-labels at a time. We are implicitly assuming, therefore, that the relative dynamics of any two spin-labels is not influenced by the presence of a third spin-label. This assumption seems plausible in this case, but it is not a pre-requisite for the EBMetaD method, which can be applied to single spin-label pairs, as mentioned. To evaluate the new methodology, three calculations were carried out: a 600-ns conventional MD simulation; a 400-ns EBMetaD simulation in which the experimental error is not considered; and a 670-ns EBMetaD simulation that does consider the confidence bands. The resulting distance distributions after equilibration are reported in **Fig. 9** (see caption for details on the analysis). Overall, the unbiased probability distributions obtained with standard MD are outside the confidence bands particularly for T4L 109/134 (**Fig. 9A**). Nonetheless, consistent with our previous work (18), the distributions calculated with EBMetaD while neglecting the error match the target with great accuracy (**Fig. 9A**). By contrast, when the confidence bands are targeted, the EBMetaD results no longer match the optimal-fit distributions and instead draw nearer the unbiased data while being fully consistent with the experimental confidence bands (**Fig. 9B**). A compelling example of this behavior is observed for the T4L 62/134 pair, for which the second peak near 45 Å in the experimentally-determined probability distribution is not fully realized in the results of the EBMetaD sampling (red lines, **Fig. 9B**), precisely because the confidence bands (black bands, **Fig. 9B**) indicate very large uncertainty in this region.

**Figure 9.**
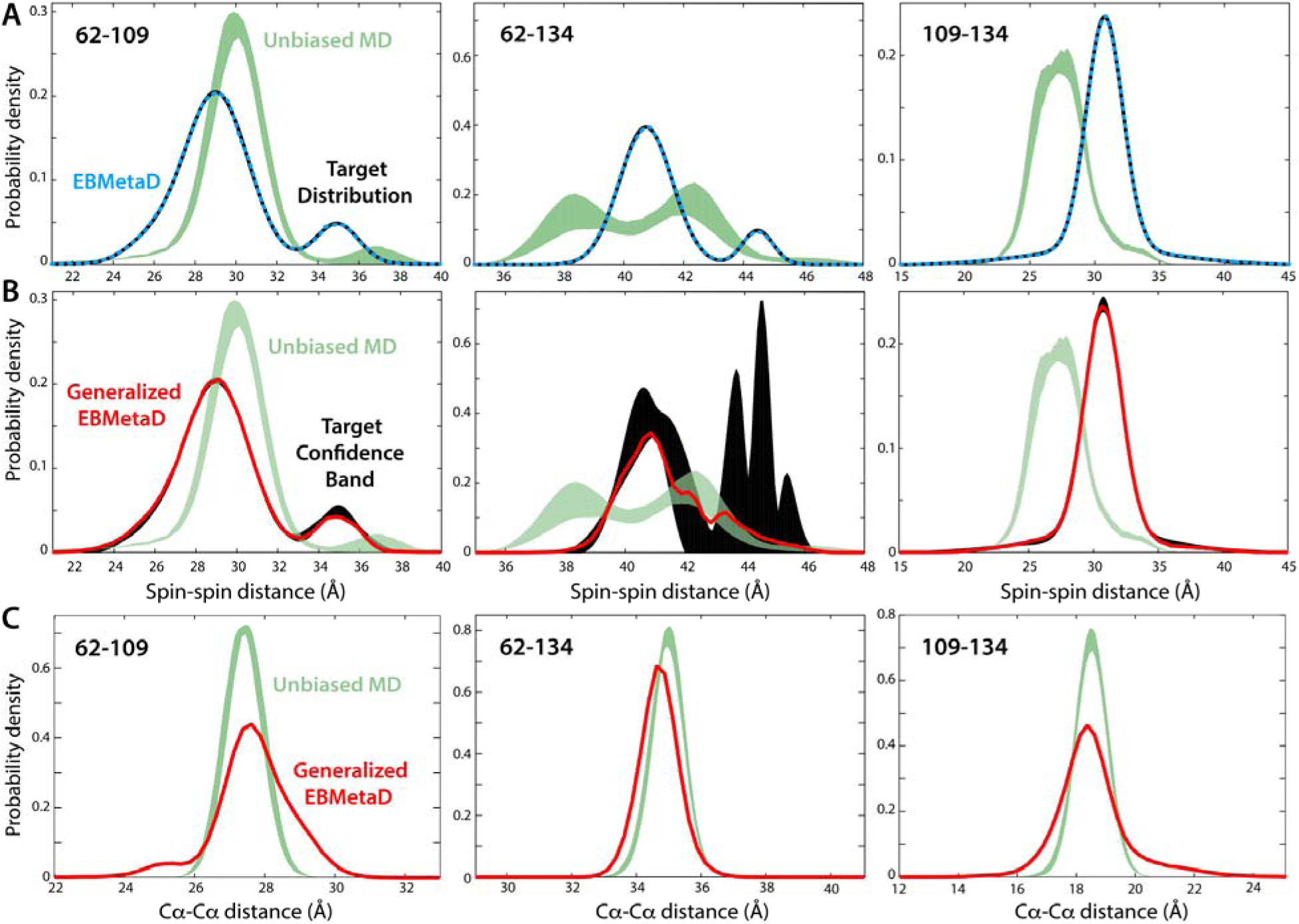
Evaluation of extended EBMetaD for spin-labeled T4 lysozyme. (A) Best-fit probability distributions obtained for each of the three experimental data sets (black) are compared with those calculated with standard MD simulations (green bands) or with EBMetaD when the experimental distribution is the target and the confidence bands are not considered (cyan). (**B**) Confidence bands from **Fig. 8** (black bands), but shown at 1σ, are compared with probability distributions calculated with EBMetaD now targeting these confidence bands (red) and the unbiased MD data. The standard MD data in panels A and B are shown as a band whose width is the standard error over 5 consecutive blocks of 100 ns each. The standard errors of EBMetaD distributions are barely visible and therefore not shown, for clarity. (**C**) Probability distributions of the distance between the Cα atoms for each of the spin-label pairs, either from standard MD (green bands) or from the EBMetaD simulations (red lines) targeting the confidence bands shown in panel (B).

Consistent with the maximum-entropy principle underlying our methodology, the magnitude of the EBMetaD biasing potential is, overall, smaller when the simulation targets the confidence bands. Specifically, the work value calculated using Eq. 28 changes from 0.96 to 0.77 kcal/mol when the confidence bands are targeted rather than the optimal-fit distributions. The small magnitude of these values reflects the limited structural dynamics of T4L which implies that the unbiased MD distributions are not entirely unlike those measured. Indeed, at the structural level, the ensemble produced by EBMetaD differs primarily from the unbiased simulation in the distribution of rotameric states of the spin-labels (**Fig. 9C, 10A**). Nevertheless, the work values derived from the biasing potential are in good agreement with those predicted from the Kullback-Leibler divergence (Eq. 30) of the unbiased and target distributions (**Fig. 9A-B**), namely 0.99 and 0.83 kcal/mol, respectively. It seems clear, therefore, that the methodology will be sufficiently sensitive to conformational changes of a larger scale.

**Figure 10.**
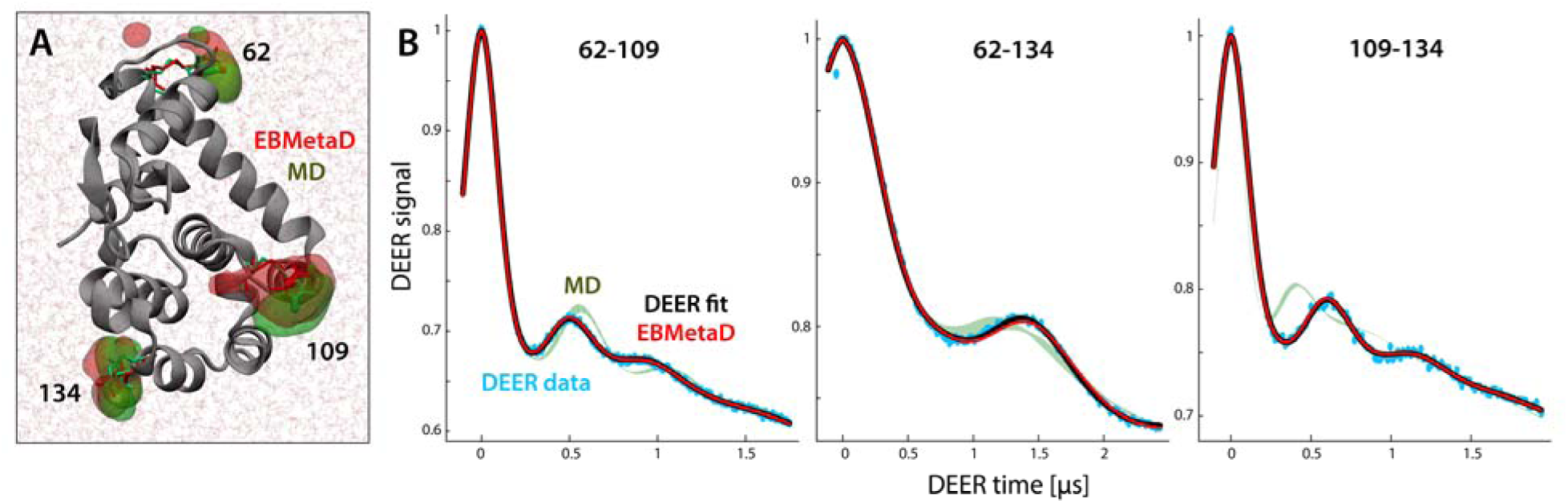
Evaluation of generalized EBMetaD for spin-labeled T4 lysozyme. (**A**) Close-up of T4 lysozyme in the MD simulation system, highlighting the spin-labels at positions 62, 109 and 134. Red and green surfaces encompass the regions occupied by the nitroxide groups during the EBMetaD and MD simulations, respectively. The RMS deviation of the protein backbone (grey cartoons) relative to the initial X-ray structure (40) is within 2 Å in both cases. (**B**) For each of the three spin-label pairs, the experimental DEER signals (cyan), and the corresponding fits (black) are compared with theoretical signals calculated (Eqs. 2-6) from the simulated EBMetaD distributions (red) in **Fig. 9B**, i.e. in consideration of the confidence bands. The calculated signals from the unbiased MD data are also shown for comparison (green bands). The Δ and *λ* parameters for each of these calculated time-traces are provided in **Supplemental Table S7**.

The molecular ensembles produced by generalized EBMetaD method not only facilitate a structural interpretation of the DEER data, but also provide a self-consistency check of the model-based analysis described above and the resulting confidence bands. That is, from each of the simulated EBMetaD ensembles, it is possible to derive the DEER time trace for each of the spin-label pairs (9, 10, 13), which can be then compared with the actual experimental signals. As the data in **Fig. 10B** demonstrates, the EBMetaD ensembles produced for each spin-label pair by targeting the confidence bands are in excellent agreement with the DEER measurements, demonstrating the consistency of the model-based analysis and the EBMetaD method. This result is particularly remarkable for T4L 62/-134, for which, as mentioned, the uncertainty band is very broad for the component at the longest distance.

In summary, atomistic simulations of T4-lysozyme demonstrate that the combined use of the model-based analysis and EBMetaD is a rigorous, self-consistent methodology to efficiently generate conformational ensembles that optimally represent DEER spectroscopic data.

## DISCUSSION

While the importance of defining experimental uncertainty and estimating errors in derived quantities is well established in science, practitioners of DEER spectroscopy have been slow to adopt methods for estimating the uncertainty in the distance distributions obtained from analyzing DEER data (13). Within the context of the model-free TR approach to the analysis of DEER data, Jeschke and coworkers (6) have developed a validation tool for estimating the uncertainty in *P*(*R*) due to variation in the background correction and the noise in the data. Edwards and Stoll have developed a Bayesian approach for estimating the uncertainty in *P*(*R*) due to the noise in the data and the regularization process (13). Alternatively, DEER data can be analyzed by modeling *P*(*R*) as a sum of Gaussian components (9, 10). The advantages of this model-based approach include the ability to analyze DEER data without the need for *a priori* background correction, the ability to perform global analysis of multiple data sets (e.g. to model functionally-relevant ligand-induced conformational changes (41-44)), and the ability to perform rigorous statistical analysis of the fit results. The major disadvantages are the need to specify particular basis functions which may deviate from the shape of the true distance distribution and the need to ensure that 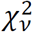 space has been fully explored to find the true global minimum.

In this work, we have extended our model-based approach to allow for a direct estimation of a confidence band for *P*(*R*) using propagation of errors, *i.e*. the delta method. By construction, this confidence band includes contribution due to both the noise in the data and the uncertainty in the background factor. The robustness of the methodology has been demonstrated through the extensive Monte Carlo analysis of simulated data. In particular, these results establish how the noise level and the maximum observed dipolar evolution time, *t*_max_, influence the uncertainty in the fit parameters and *P*(*R*). From the results in **Fig. 5** and **Supplemental Tables S4** and **S5** it is evident that precise estimates of the contribution of a given component to *P*(*R*) can be obtained as long as *one* full modulation period can be observed within *t*_max_, *i.e*. 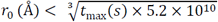. This equation is similar to one proposed by Jeschke for the minimum *t*max required to determine a *r*_0_ (mean) value (3). Even under less than optimal conditions, with extremely short *t*_max_, reasonable estimates of *P*(*R*) can be obtained (see **Fig. 4** and **Supplemental Tables S2)**. It should be noted that different analysis methods can influence the reliability of the determined *P*(*R*) as much as the information content in the data itself.

In conjunction with our approach for determining confidence bands on distance distributions obtained from fitting DEER data, we have also proposed an atomistic simulation method that imposes a minimum-information bias on a molecular dynamics trajectory so that the conformational ensemble explored reproduces one or more experimental probability distributions as precisely as dictated by their confidence bands. This technique is an extension of the Ensemble-Biased Metadynamics (EBMetaD) method (18) reformulated here to include the confidence bands in the target distributions. Although other enhanced-sampling approaches exist that use probability distributions as input data (17, 18), to our knowledge this is the first report of a methodology that also considers the uncertainties in those distributions.

The proposed methodology is evaluated for T4L in explicit water, simultaneously targeting the distance distributions obtained from three spin-labeled double mutants. It is clearly shown how the EBMetaD ensembles deviate from those obtained from conventional MD simulations precisely so that the calculated probability distributions draw inside the experimental confidence bands. For values of *R* for which the experimental uncertainty of *P*(*R*) is small, the simulation approaches the best-fit distributions as accurately as required by their confidence bands. In poorly defined regions, EBMetaD behaves comparably to an unbiased MD simulation. Needless to say, MD trajectories are inexact on account of the many approximations and simplifications inherent to this technique Thus, while the EBMetaD method guarantees that the simulated ensemble will reproduce the input experimental data, it does not guarantee that this ensemble is free of error otherwise, or that it represents the only possible solution. It is also worth noting that the EBMetaD method is not limited to the interpretation of DEER data. Indeed, this approach may be used for any observable for which a probability distribution can be derived experimentally (or postulated theoretically), as long as it is computable during run-time from the set of atomic coordinates in the molecular system.

The EBMetaD simulation methodology is based on the fundamental concept of maximum entropy. Thus, the simulation uses the minimum information required to reproduce the target data and therefore the bias introduced to modify the simulated ensemble is also minimal. This notion becomes clear when the magnitude of the bias applied is transformed into a quasi-equilibrium work value. Specifically, the smaller the uncertainty, the greater the amount of work required to reproduce a given experimental distribution. It is worth pointing out that not all molecular refinement methods satisfy this intuitive minimum-information principle. Standard procedures to construct structural ensembles from NMR data, for example, impose distance restraints on pairs of atoms that imply specific distributions around the target value (45). In doing so, these computational approaches utilize more information than what the experiment actually provides, which is only the ensemble-average value of those distances and not their distribution. We therefore anticipate that variations of this maximum-entropy paradigm, together with high-end molecular-simulation methods, will become increasingly used to derive a rigorous interpretation of a variety of spectroscopic data.

The possibility of quantifying the minimum work required to generate a conformational ensemble that is most consistent with a given experimental probability distribution is a notable feature of the EBMetaD approach also from a mechanistic standpoint. It is not uncommon that different structures are known for a biomolecule, often presumed to represent distinct and interconverting functional states. Using DEER spectroscopy under different experimental conditions, different components of *P*(*R*) can be assigned to these different functional states. EBMetaD simulations provide a means to relate structural and spectroscopic data: the experimental structure that, when simulated, requires the least amount of work to reproduce a given set of DEER data can be assumed to be the best representative of the conditions used to collect that spectroscopic signal. It should be noted, though, that the distance between two or more spin-labels might not be an appropriate reaction coordinate to drive the reversible exploration of large-scale or intricate conformational changes. In such cases, the EBMetaD approach ought to be combined with other strategies devised to enhance the sampling of those conformational changes. Multiple-walker algorithms such as Bias-Exchange Metadynamics (46, 47) would be a natural choice to integrate EBMetaD with other biasing schemes, but other options are also possible.

While there are compelling reasons to interpret the *P*(*R*) obtained in terms of components corresponding to distinct functionally relevant structures. There are a number of factors that can give rise to artifactual components in the *P*(*R*) that do not, in fact, correspond to distinct structural states. These factors include imperfect background correction and orientation selection effects. The TR approach biases the *P*(*R*) obtained to have an equal degree of smoothness across all *R*. Thus, in situations where the true *P*(*R*) may contain a mix of narrow and broad components, TR will split a single broad component into a sum of multiple narrow components (see **Supplemental Fig. S5D**). As a result, an approach that first fits the time-domain data using TR and then fits the *P*(*R*) obtained to a sum of Gaussians can overestimate the number of components.

Instead, our approach directly fits the time-domain data using a sum of components of varying width and uses *BIC* values to select the optimal number of components based on the principal of parsimony. To lend credence to the structural and functional relevance of these components, a global analysis of multiple DEER signals may reveal how the amplitudes of these components change with conditions that can be manipulated experimentally. Otherwise, care must be taken when assigning structural and biological relevance to terms in the mathematical equation used to model *P*(*R*). When using a model-based approach, particularly for high signal-to-noise data, additional terms may be required for an optimal fit to account for deviations from Gaussian shape. For example, each of the T4L data sets can be fit using one fewer component (see Supplemental Table S8 and Supplemental Figure S6) using alternate basis functions (see Supplemental Methods). While these non-Gaussian functions yield very similar fits with comparable 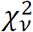 values, they are favored by *BIC* by virtue of their lower number of fit parameters. The differences in the distributions, obtained using different basis functions or TR that are shown in Supplemental Figure S6, highlight the inherent uncertainty in the shape of *P*(*R*) regardless of the method used. Ultimately, molecular simulations based on the EBMetaD method can clarify the interpretation of multicomponent distance distributions inferred from DEER experiments in terms of distinct molecular conformations, while taking into account rigorously determined uncertainties in the experimental results.

## CONCLUSIONS

In this study, we propose two complementary computational strategies that, taken together, provide a compelling methodology to derive an objective structural interpretation of DEER measurements for a dynamic biomolecular system. Ultimately, both methodologies underscore the importance of a rigorous error estimate for a correct interpretation of the experimental data.

## AUTHOR CONTRIBUTIONS

The research on the analysis of DEER data was designed by HSM, RAS, and EJH (Fig. 1-5 and 10) and then developed and implemented by EJH with the assistance of RAS. The research on EBMetaD (Fig. 6-10) was designed by FM and JDFG and developed and implemented by FM. EJH, FM, and JDFG wrote the manuscript with the assistance of HSM and RAS.

## ACKNOWLEDGEMENTS

F.M. and J.D.F. G. were funded by the Division of Intramural Research of the National Heart, Lung and Blood Institute, National Institutes of Health. HSM received funding from NIH GM 077657. EJH and RAS thank the Biostatistics Clinics service of the Vanderbilt University Department of Biostatistics for helpful discussion about the delta method.

## REFERENCES

1. Borbat, P. P., and J. H. Freed. 2014. Pulse Dipolar Electron Spin Resonance: Distance Measurements. In Structural Information from Spin-Labels and Intrinsic Paramagnetic Centres in the Biosciences. C. R. Timmel, and J. R. Harmer, editors. 1–82.

2. Jeschke, G. 2012. DEER Distance Measurements on Proteins. In Annual Review of Physical Chemistry, Vol 63. M. A. Johnson, and T. J. Martinez, editors. 419–446.

3. Jeschke, G. 2014. Interpretation of Dipolar EPR Data in Terms of Protein Structure. In Structural Information from Spin-Labels and Intrinsic Paramagnetic Centres in the Biosciences. C. R. Timmel, and J. R. Harmer, editors. 83–120.

4. McHaourab, Hassane S., P. R. Steed, and K. Kazmier. 2011. Toward the Fourth Dimension of Membrane Protein Structure: Insight into Dynamics from Spin-Labeling EPR Spectroscopy. Structure 19: 1549–1561.

5. Bowman, M. K., A. G. Maryasov, N. Kim, and V. J. DeRose. 2004. Visualization of distance distribution from pulsed double electron-electron resonance data. Applied Magnetic Resonance 26: 23–39.

6. Jeschke, G., V. Chechik, P. Ionita, A. Godt, H. Zimmermann, J. Banham, C. R. Timmel, D. Hilger, and H. Jung. 2006. DeerAnalysis2006 - a comprehensive software package for analyzing pulsed ELDOR data. Applied Magnetic Resonance 30: 473–498.

7. Chiang, Y. W., P. P. Borbat, and J. H. Freed. 2005. The determination of pair distance distributions by pulsed ESR using Tikhonov regularization. Journal of Magnetic Resonance 172: 279–295.

8. Jeschke, G., G. Panek, A. Godt, A. Bender, and H. Paulsen. 2004. Data analysis procedures for pulse ELDOR measurements of broad distance distributions. Applied Magnetic Resonance 26: 223–244.

9. Brandon, S., A. H. Beth, and E. J. Hustedt. 2012. The global analysis of DEER data. Journal of Magnetic Resonance 218: 93–104.

10. Stein, R. A., A. H. Beth, and E. J. Hustedt. 2015. A Straightforward Approach to the Analysis of Double Electron-Electron Resonance Data. Methods in enzymology 563: 531–567.

11. Fajer, P., L. Brown, and L. Song. 2007. Practical Pulsed Dipolar ESR (DEER). In ESR Spectroscopy in Membrane Biophysics. M. A. Hemminga, and L. J. Berliner, editors. Springer US. 95–128.

12. Pannier, M., V. Schädler, M. Schöps, U. Wiesner, G. Jeschke, and H. W. Spiess. 2000. Determination of Ion Cluster Sizes and Cluster-to-Cluster Distances in Ionomers by Four-Pulse Double Electron Electron Resonance Spectroscopy. Macromolecules 33: 7812–7818.

13. Edwards, T. H., and S. Stoll. 2016. A Bayesian approach to quantifying uncertainty from experimental noise in DEER spectroscopy. Journal of Magnetic Resonance 270: 87–97.

14. Tellinghuisen, J. 2001. Statistical Error Propagation. The Journal of Physical Chemistry A 105: 3917–3921.

15. Ver Hoef, J. M. 2012. Who Invented the Delta Method? American Statistician 66: 124–127.

16. Casella, G., and R. L. Berger. 2002. Statistical Inference. Duxbury Press, Pacific Grove CA.

17. Roux, B., and S. M. Islam. 2013. Restrained-ensemble molecular dynamics simulations based on distance histograms from double electron-electron resonance spectroscopy. The journal of physical chemistry. B 117: 4733–4739.

18. Marinelli, F., and José D. Faraldo-Gómez. 2015. Ensemble-Biased Metadynamics: A Molecular Simulation Method to Sample Experimental Distributions. Biophysical Journal 108: 2779–2782.

19. Polyhach, Y., E. Bordignon, and G. Jeschke. 2011. Rotamer libraries of spin labelled cysteines for protein studies. Physical Chemistry Chemical Physics 13: 2356–2366.

20. Fajer, P., M. Fajer, M. Zawrotny, and W. Yang. 2015. Simulation of spin label structure and its implication in molecular characterization. Methods in enzymology 563: 623–642.

21. Kazmier, K., N. S. Alexander, J. Meiler, and H. S. Mchaourab. 2011. Algorithm for selection of optimized EPR distance restraints for de novo protein structure determination. J Struct Biol 173: 549–557.

22. Islam, S. M., R. A. Stein, H. S. McHaourab, and B. Roux. 2013. Structural Refinement from Restrained-Ensemble Simulations Based on EPR/DEER Data: Application to T4 Lysozyme. The Journal of Physical Chemistry B 117: 4740–4754.

23. Milov, A. D., A. G. Maryasov, and Y. D. Tsvetkov. 1998. Pulsed electron double resonance (PELDOR) and its applications in free-radicals research. Applied Magnetic Resonance 15: 107–143.

24. Burnham, K. P., and D. R. Anderson. 2002. Model selection and multimodel inference: a practical information-theoretic approach Springer-Verlag, New York.

25. Edwards, T. H., and S. Stoll. 2018. Optimal Tikhonov regularization for DEER spectroscopy. Journal of magnetic resonance (San Diego, Calif.: 1997) 288: 58–68.

26. Bevington, P. R., and D. K. Robinson. 1992. Data reduction and error analysis for the physical sciences. McGraw-Hill, New York.

27. Press, W. H., S. A. Teukolsky, W. T. Vetterling, and B. P. Flannery. 1993. Numerical Recipes in FORTRAN; The Art of Scientific Computing. Cambridge University Press.

28. Phillips, J. C., R. Braun, W. Wang, J. Gumbart, E. Tajkhorshid, E. Villa, C. Chipot, R. D. Skeel, L. Kale, and K. Schulten. 2005. Scalable molecular dynamics with NAMD. J Comput Chem 26: 1781–1802.

29. MacKerell, A. D., D. Bashford, M. Bellott, R. L. Dunbrack, J. D. Evanseck, M. J. Field, S. Fischer, J. Gao, H. Guo, S. Ha, D. Joseph-McCarthy, L. Kuchnir, K. Kuczera, F. T. K. Lau, C. Mattos, S. Michnick, T. Ngo, D. T. Nguyen, B. Prodhom, W. E. Reiher, B. Roux, M. Schlenkrich, J. C. Smith, R. Stote, J. Straub, M. Watanabe, J. Wiorkiewicz-Kuczera, D. Yin, and M. Karplus. 1998. All-atom empirical potential for molecular modeling and dynamics studies of proteins. Journal of Physical Chemistry B 102: 3586–3616.

30. Mackerell, A. D., Jr., M. Feig, and C. L. Brooks, 3rd. 2004. Extending the treatment of backbone energetics in protein force fields: limitations of gas-phase quantum mechanics in reproducing protein conformational distributions in molecular dynamics simulations. J Comput Chem 25: 1400–1415.

31. Sezer, D., J. H. Freed, and B. Roux. 2008. Parametrization, molecular dynamics simulation, and calculation of electron spin resonance spectra of a nitroxide spin label on a polyalanine alpha-helix. Journal of Physical Chemistry B 112: 5755–5767.

32. Roux, B., and J. Weare. 2013. On the statistical equivalence of restrained-ensemble simulations with the maximum entropy method. Journal of Chemical Physics 138.

33. Marinelli, F., and G. Fiorin. 2018. Structural characterization of biomolecules through atomistic simulations guided by DEER measurements. Under Review.

34. Cesari, A., A. Gil-Ley, and G. Bussi. 2016. Combining Simulations and Solution Experiments as a Paradigm for RNA Force Field Refinement. Journal of Chemical Theory and Computation 12: 6192–6200.

35. Hummer, G., and J. Kofinger. 2015. Bayesian ensemble refinement by replica simulations and reweighting. J Chem Phys 143: 243150.

36. Wang, W., and M. A. Carreira-Perpiñán. 2013. Projection onto the probability simplex: An efficient algorithm with a simple proof, and an application. arXiv:1309.1541 [cs.LG].

37. Plimpton, S. 1995. Fast Parallel Algorithms for Short-Range Molecular-Dynamics. J Comput Phys 117: 1–19.

38. Fiorin, G., M. L. Klein, and J. Hénin. 2013. Using collective variables to drive molecular dynamics simulations. Molecular Physics 111: 3345–3362.

39. Crespo, Y., F. Marinelli, F. Pietrucci, and A. Laio. 2010. Metadynamics convergence law in a multidimensional system. Phys Rev E Stat Nonlin Soft Matter Phys 81: 055701.

40. Weaver, L. H., and B. W. Matthews. 1987. Structure of bacteriophage T4 lysozyme refined at 1.7 A resolution. J Mol Biol 193: 189–199.

41. Collauto, A., H. A. DeBerg, R. Kaufmann, W. N. Zagotta, S. Stoll, and D. Goldfarb. 2017. Rates and equilibrium constants of the ligand-induced conformational transition of an HCN ion channel protein domain determined by DEER spectroscopy. Physical Chemistry Chemical Physics 19: 15324–15334.

42. Kazmier, K., S. Sharma, M. Quick, S. M. Islam, B. Roux, H. Weinstein, J. A. Javitch, and H. S. McHaourab. 2014. Conformational dynamics of ligand-dependent alternating access in LeuT. Nature structural & molecular biology 21: 472–479.

43. Martens, C., R. A. Stein, M. Masureel, A. Roth, S. Mishra, R. Dawaliby, A. Konijnenberg, F. Sobott, C. Govaerts, and H. S. McHaourab. 2016. Lipids modulate the conformational dynamics of a secondary multidrug transporter. Nature structural & molecular biology 23: 744–751.

44. Mishra, S., B. Verhalen, R. A. Stein, P. C. Wen, E. Tajkhorshid, and H. S. McHaourab. 2014. Conformational dynamics of the nucleotide binding domains and the power stroke of a heterodimeric ABC transporter. eLife 3:e02740.

45. Schwieters, C. D., J. J. Kuszewski, and G. M. Clore. 2006. Using Xplor-NIH for NMR molecular structure determination. Prog Nucl Mag Res Sp 48: 47–62.

46. Marinelli, F., S. I. Kuhlmann, E. Grell, H. J. Kunte, C. Ziegler, and J. D. Faraldo-Gomez. 2011. Evidence for an allosteric mechanism of substrate release from membrane-transporter accessory binding proteins. P Natl Acad Sci USA 108:E1285–E1292.

47. Liao, J., F. Marinelli, C. Lee, Y. H. Huang, J. D. Faraldo-Gomez, and Y. X. Jiang. 2016. Mechanism of extracellular ion exchange and binding-site occlusion in a sodium/calcium exchanger. Nature structural & molecular biology 23: 590–599.

